# Cytokine Regulation of Human Antibody Responses to Influenza Vaccines

**DOI:** 10.1101/2025.08.07.668218

**Authors:** Guangbo (Bill) Chen, Jing Guo, John Heath, Tyler Prestwood, Woo Joo Kwon, Ashley Smith, Tran T. Nguyen, Elsa Sola, AbuBakr Sangare, Vamsee Mallajosyula, Ryan Ichiro Furuichi Fong, Azam Mohsin, Lei Chen, Oviya Siva, Cindy Padilla, Mingdian Tan, Cornelia L. Dekker, Philip Grant, Ying Lu, Harry B. Greenberg, William H. Robinson, Catherine Blish, Shai Shen-Orr, Ahmad Salehi, Holden T. Maecker, Purvesh Khatri, Paul J. Utz, Yueh-hsiu Chien, Mark M. Davis

## Abstract

Studies show that human vaccine responses widely vary. Here, we analyzed data measuring 66 cytokines from 4 different inactivated influenza vaccine (IIV) cohorts over 5 seasons (N=581) and identified a significant correlation between baseline/day 0 serum IL-18 and IFN-β concentrations and the vaccine-specific antibody response on day 28, suggesting these factors may have an adjuvant-like effect. To investigate this further, we tested the impact of 19 cytokines on the development of anti-influenza antibody response when administered together with the IIV vaccine in human tonsil and spleen organoids. We found that Type I IFNs (IFN-β and others), IL-21, IL-12, IL-10, but not IL-18 or IFN-γ, enhanced the antibody response. The live attenuated influenza vaccine (LAIV) induced a stronger antibody response than the inactivated one in organoids. Adding a single cytokine, IFN-β, to IIV stimulation recapitulated most of the live vaccine-specific cytokine activation program. It increased the antibody response of the inactivated vaccine to that of the LAIV. Two other antibody-boosting cytokines, IL-12 and IL-21, were induced by LAIV but not by Type I IFNs, indicating different cytokines can affect different pathways leading to a robust antibody response. We then generated cytokine mRNA lipid nanoparticles (LNPs) to test the effect of cytokines on IIV response in a mouse model of immunization. We found that IL-21-LNPs augmented the quantity and breadth of the antibody responses, while IFN-β LNPs enhanced durability. These findings identified parallel cytokine pathways regulating human vaccine responses and provide a rationale for using cytokines as adjuvants to mimic the effectiveness of live-attenuated vaccines without the risk of viral replication.

## Background

Each year, approximately 800 million people receive influenza vaccine, with 98% receiving the inactivated influenza vaccine (IIV), The elicited specific antibody response (associated with the protection against infection) vary widely and the overall protection rate ranges from 20% and 60%, depending on the prevalent strains, with most morbidity and mortality occurring in older individuals (> 65 years)^1^. Understanding the immunological factors that differentiate between strong and weak responders can inform us of the potential mechanisms driving vaccine responses and suggest ways to improve them.

In our previous investigation of human cohorts, we found that while the specificity of the response after influenza vaccination was determined by host genetics^2^, the variability in the magnitude of this response was mostly driven by non-heritable factors^3^. Such factors may have included prior cytomegalovirus infection and gut microbiota (e.g., activating TLR5^5,6^). Immune states at baseline or early vaccine responses may correlate and potentially regulate the vaccine response^7–9^. Multiple comprehensive studies conducted over the past decade have characterized the pre-vaccination or early post-vaccination immune transcriptome signatures that correlated with improved antibody responses to the influenza vaccine^10–15^. Several of these transcriptome signatures contained interferon (IFN)-inducible genes^10–13,16^; however, it has been unclear whether Type I or Type II interferons were responsible for these signatures and whether they were correlative or causal.

Here, we reported the identification of multiple cytokines that correlated with elevated antibody responses in human influenza virus vaccination studies. We combined data from four influenza vaccine cohorts in five flu seasons (N=581). We found a robust correlation between the pre-vaccination (pre-VAX) serum abundance of cytokines and post-VAX flu antibody responses in a subpopulation with a lower baseline antibody titer (N=389). To distinguish between cytokines that merely correlate with an enhanced response to a vaccine versus those that could drive the response, we modified a human immune organoid system we initially developed for tonsil organoids^20^ to use spleen cells derived from organ donors, using a protocol developed by Kathuria et al. (submitted) and using a 96-well format^21^. The platform used much less cell input (97% lower). It enabled high-throughput screening and identified several cytokines that can enhance the antibody response elicited by the IIV in humans. Lastly, we wanted to test whether these in vitro results would hold up in an in vivo system and since RNA vaccines are known to be transported to lymph nodes^22^, we created mRNA-LNP constructs encoding some of these cytokines and tested them in a mouse model. While the mRNA-LNP itself had a significant effect^23^, constructs carrying IL-21 and IFN-β enhanced this even further, showing that some cytokines could be effective adjuvants.

## Results

### Baseline antibody titer influences the vaccine response phenotype

Given the central role of cytokines in modulating immune responses^24,25^, we analyzed specific pre-vaccination (pre-VAX) serum cytokine profiles and the magnitude of the post-VAX antibody response (**Fig. 1a**). To achieve sufficient statistical power, we pooled data from four different flu vaccine cohorts from five different seasons (**Fig. 1b**). In these studies, inactivated influenza vaccine (IIV) was administered to subjects, and serum was analyzed in pre-VAX (day 0) and post-VAX (day 28) time points. Influenza (flu) hemagglutinin inhibition (HAI) antibodies for the vaccinated strains were determined at both time points, with the ratio defined as the antibody response. Considerable heterogeneity was present among cohorts in age, sex, flu strains, dose included in the vaccine, and cytokine profiling technology in different seasons.

**Fig. 1:**
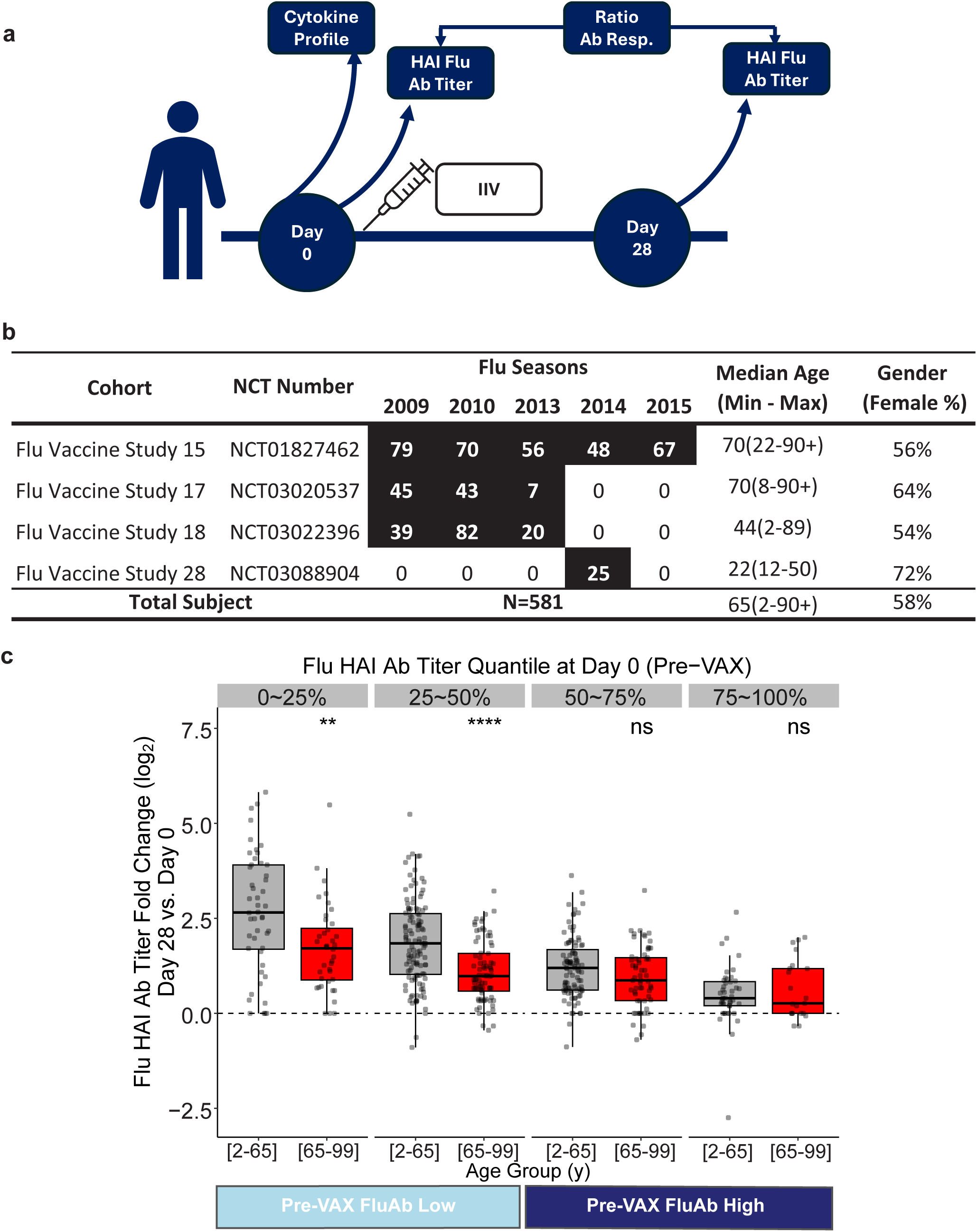
Age effects on vaccine responses were only detectable in the subjects with low pre-vaccination flu antibodies. a) The scheme of the human cohort studies investigating the correlation between serum cytokine abundance and antibody response (ratio of HAI flu antibody abundance between Day 28 and Day 0). **b)** The summary statistics of samples and cohorts included in the study. **c)** The correlation between Pre-VAX flu hemagglutinin inhibition (HAI) antibody titers and Post-VAX antibody response. The antibody response was defined by the log2 ratio of the Day 28 and Day 0 flu HAI antibody titers. All subjects who received a lower dose (15ug/vaccine strain) IIV were included (while the older adults (>65 y) received the high-dose (60ug/vaccine strain) formulation). Subjects were stratified into quartiles based on pre-vaccination (Day 0) HAI titers. Pre-VAX low and high referred to the lower and upper 50% of pre-vaccination HAI titers, respectively; most in the low group had titers <40, a non-protective threshold. We performed Wilcoxon ranking tests between different age groups. **, p<0.01; ***, p<0.001.

The correlation between pre-VAX (Day 0) flu antibody titer and post-VAX (Day 28) antibody response also showed great heterogeneity (**Fig. 1c**). To address this, we divided the population into 4 quartiles based on their pre-VAX flu antibody titers. In line with prior study, we observed that both the median and dynamic ranges of antibody responses decreased with an increasing abundance of pre-VAX antibodies, potentially due to the antigen clearance and thus lower vaccine “take”^10,26,27^. Aging is known to lower vaccine response significantly. Still, interestingly, we found that responses were only lower in older individuals for subjects with low pre-VAX flu antibodies (quantile 1 and 2, lower 50 percentile, **Fig. 1c**). These results suggested that the correlation between vaccine response and a biological characteristic (such as age) was considerably stronger in people with a low level of pre-VAX flu antibodies (and potentially higher vaccine “take”). Compared with the group with a high level of pre-VAX flu antibodies, the low-level group had a higher risk of flu infection{Fox, 1982 #3;Ng, 2013 #4}. Among the 389 subjects with a pre-VAX flu HAI antibody titer lower than 40 (deemed not protected), 89% of them were in the group with a low level of pre-VAX flu antibodies. For this reason, our analysis focused on this group.

### Meta-analysis identifies a correlation between pre-VAX cytokine abundance and post-VAX antibody response to the flu vaccine

Heterogeneity (due to technical and biological factors) was a major challenge for pooling data from different cohorts. Here, we divided the dataset by confounders (age, vaccine dose, sex, day 0 flu antibody quantile, cohort, flu season) into sub-populations and performed correlations with each sub-population (see an example of the grouping in **Fig. S5a**). We then summarized the results using meta-analysis, a method capable of identifying robust signals among heterogeneous data^29^.

We performed a meta-analysis of 66 cytokines measured across the four cohorts. The analysis identified 2 cytokines (IL-18 and IFN-β) and 1 chemokine (GRO-α, also known as CXCL1) whose pre-VAX serum abundance significantly correlated with post-VAX antibody response (False Discovery Rate <5%, **Fig. 2**). IFN-β and IL-18 activate two different immune signaling pathways. IFN-β is a type I interferon (IFN) that is critical for antiviral immunity, including antibody response^30^. Among the three members of the type I IFNs tested (IFN-α, IFN-β, IFN-ω), IFN-β has the highest binding affinity to the Type I IFN receptor (IFNAR1/2), which is expressed by many cell types, including B cells^31^. Interleukin-18 (IL-18) is a multifaceted cytokine primarily known for its role in stimulating T cells to produce the Type II interferon (IFN-γ)^32,33^. We also identified several cytokines with a borderline significant correlation (FDR <10%), which included T Helper cytokines IL-17 and IL-9 (**Fig. 2**).

**Fig. 2:**
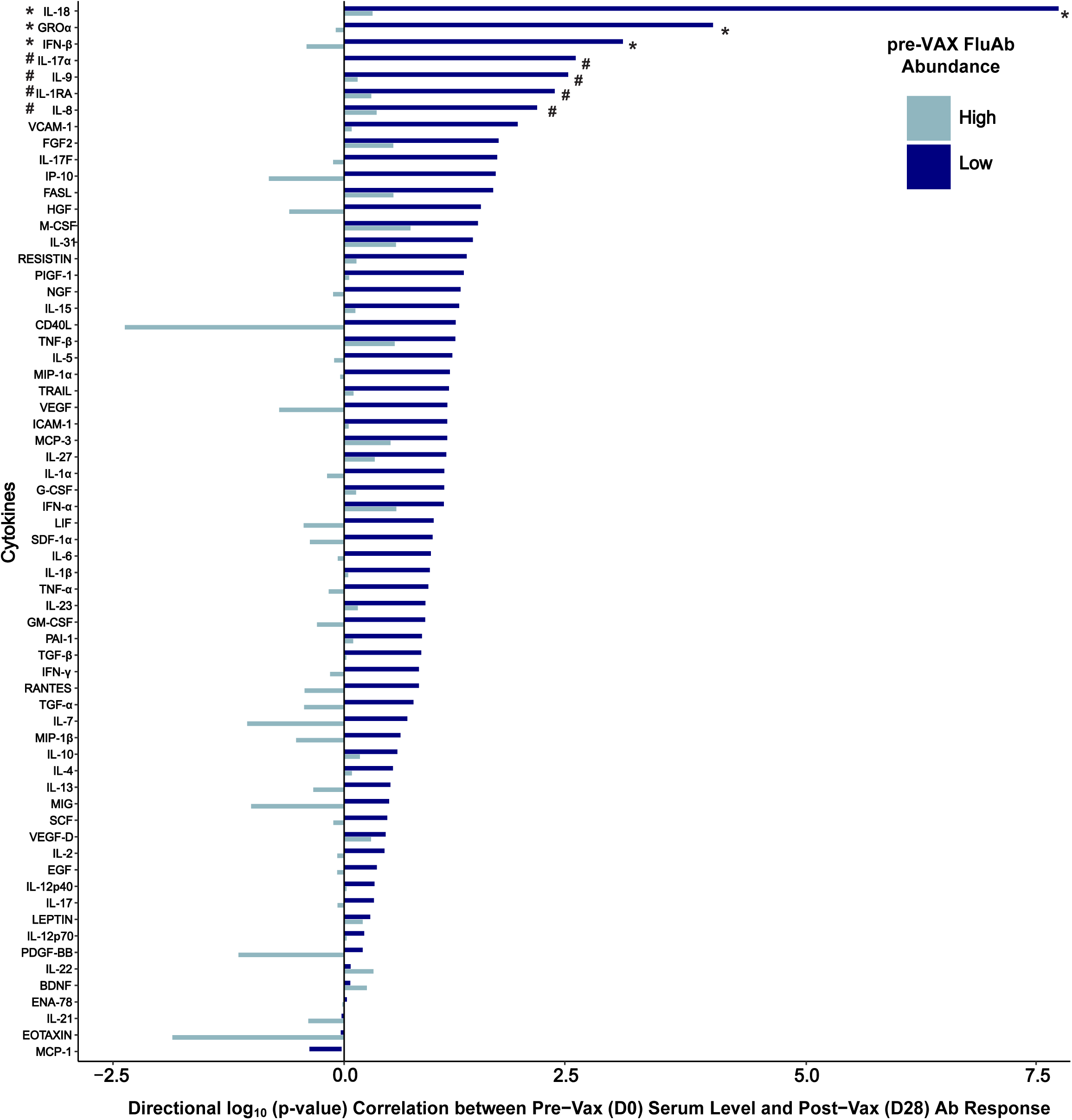
Meta-analysis identified a correlation between pre-VAX serum abundance of cytokines and post-VAX antibody response against the flu vaccine (IIV). We examined the correlation for 66 cytokines across the cohorts included in Fig. 1b. The correlations were performed separately in groups with low or high Pre-VAX flu HAI antibody titers, as defined in Fig. 1c. Details of the meta-analysis of the correlation can be found in the supplementary information. Log10 P values of the correlations were provided with the sign denoting the direction of the correlations (positive or negative slopes). False discovery rates were reported for significant or borderline significant hits. Details about the correlation and meta-analysis can be found in the supplementary methods.

### Functional screen in human spleen organoids identifies cytokine adjuvants, including Type I IFNs but not Type II IFN

Human cohort studies can identify correlations of immune function, but they cannot prove causality without interventional studies. While the functionality of many cytokines has been tested in various mouse vaccination models, the relevance to the human vaccination process has not been examined. Moreover, each mouse model typically tested only 1 or 2 cytokines. When different cytokine adjuvant effects were compared across different mouse studies, there was heterogeneity in cytokine delivery formats (DNA, protein), antigen, and delivery route used (intramuscular, intranasal, intraperitoneal, etc., **Table S1 in Supplementary Info.**). A functional systems immunology approach that compares different molecules delivered in the same format and assayed against the same antigen in a human culture system can allow quantitative evaluation of adjuvant effects and prioritize the adjuvant candidates. Recently, we developed a human immune organoid system enabling efficient hypothesis testing. The immune organoids are primary culture, different from many other organoids derived from tissue stem cells or induced Pluripotent Stem Cells (iPSCs). The immune organoids retain memory T and B cells from the donors, including those carrying memory against the influenza infections/vaccines. Meanwhile, the immune organoids recapitulate key antibody response features, including the production of antigen-specific antibodies, somatic hypermutation, and affinity maturation^21,34^. Building on our prior work, we devised a low-cell-input immune organoid culture system (decreased the cell input number by 97% to 1.6 × 10^5 cells), which allowed us to generate many immune organoid cultures in a 96-well format. We performed a screen and systematically examined cytokines’ adjuvant functionality to the inactivated vaccine (**Fig. 3a, a functional systems immunology approach**). We harvested 3 tonsils from patients from the clinic. In collaboration with the Donor Network West, an Organ Processing Organization (OPO), that serves Northern California and Northern Nevada, we procured spleens from 2 authorized ventilated deceased donors. Using this higher throughput immune organoid format, we generated 5 sets of immune organoids from 5 different subjects. We administered them with IIV, alone, or in combination with 19 different cytokines. Most of these cytokines have been shown to boost antibody response in one or multiple mouse vaccination models (**Table S1**). We combined the data since the cytokines’ adjuvant effects are similar between spleen and tonsil organoids (**Fig. S1**). For each cytokine, we tested three different concentrations, ranging from 1 ng/mL to 100 ng/mL. However, for IL1-β (10 times lower at each concentration), and IL-18 (10 times higher at each concentration), we used different concentrations as their physiological concentrations in human serum differ from those of other cytokines^35^. Antibody production (relative to the IIV-only controls) on day 7 (**Fig. 3a**) was measured. In total, we assayed 58 different conditions, including the control, across five biological replicates. The screen revealed that all type I IFNs (IFN-β and IFN-ω, though IFN-α was less effective) enhanced antibody production induced by the inactivated vaccine (**Fig. 3b**). Moreover, we discovered potent adjuvant function of several other cytokines. Among them, IL-21 and IL-9, which are secreted by T follicular helper and Th9 cells, respectively^36^. IL-12 induces the formation of Th1 and Tfh cells^37^. IL-10 enhances B cell survival and proliferation^38^. IL-10 can also act as one of the downstream effectors of type I IFN (also see below). For most factors, the effect was detected at concentrations as low as 1ng/ml, at the picomolar level. Notably, IL-18 or its downstream effector, type II IFN (IFN-γ), did not enhance the IIV-induced antibody response in human spleen and tonsil organoids (**Fig. 3b**), which is in contrast to the strong correlation observed between IL-18 abundance and antibody response in human cohorts.

**Fig. 3:**
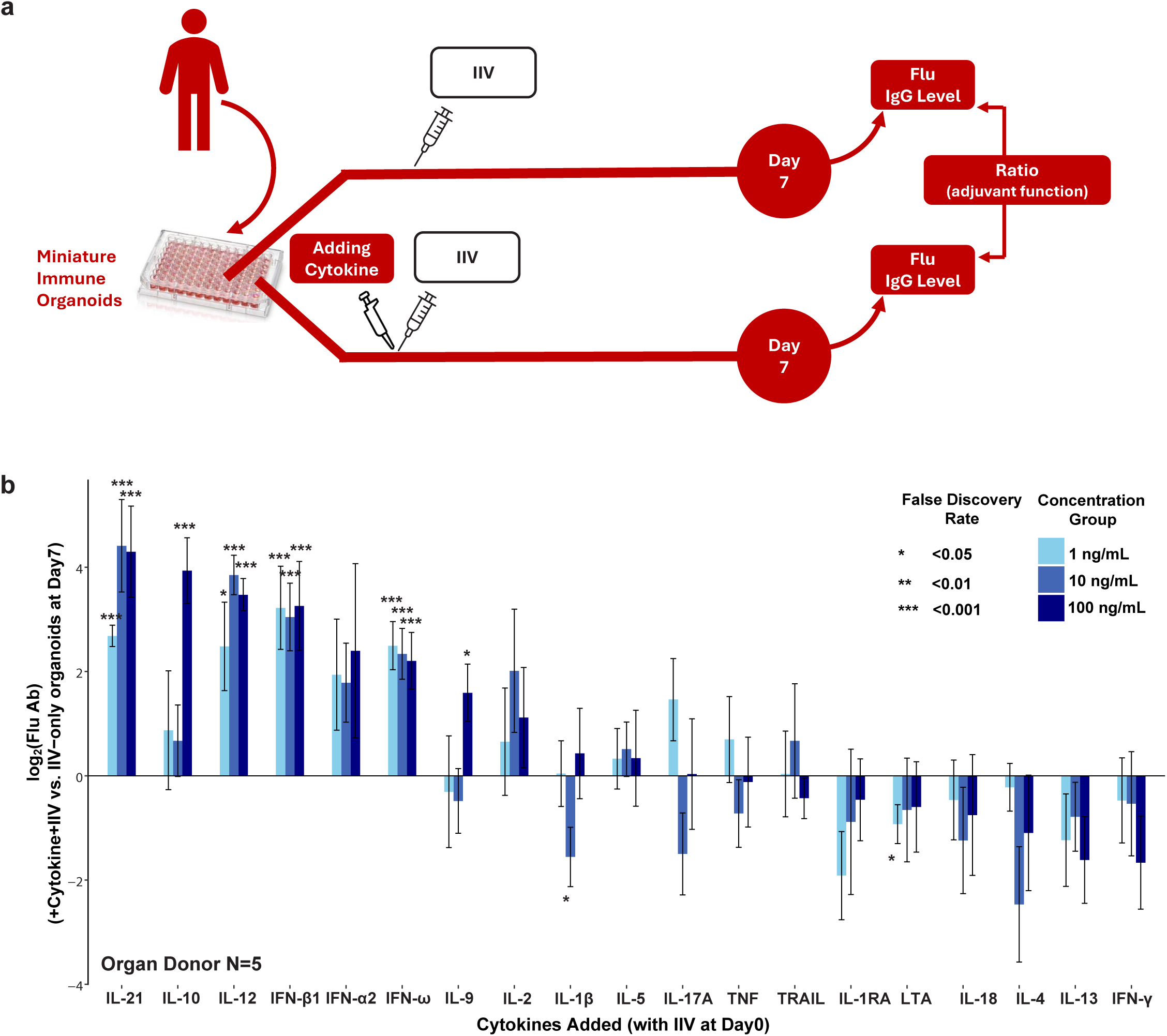
Functional screening in human immune organoids identified IIV-adjuvating cytokines. **a)** We tested the cytokine adjuvant function in 3 tonsil- and 2 spleen-organoids. We added 19 different cytokines together with the vaccine (IIV), and for each cytokine, we tested 3 different concentrations ranging from 1 ng/ml to 100 ng/ml, except IL1β (10x lower for each concentration) and IL18 (10x higher for each concentration). We measured the antibody production (relative to non-cytokine-added IIV only controls) on Day 7. As the results from the tonsil and spleen organoids are similar (**Fig. S1**), they are combined. **b)** The assay results are presented.

### Type I IFNs drive an immune activation program that mimics a live vaccine

While cytokines can regulate vaccine response, vaccination can also induce cytokines that coordinate the antibody response^39–42^. We exposed spleen organoids to IIV, and collected the supernatant on Day 3, analyzing the samples using a DNA-barcoded multiplexed cytokine detection technology with femtomolar sensitivity (NULISA)^43^ (**Fig. 4a**). Of the 250 cytokines/chemokines assayed, 40 were significantly induced by IIV treatment (**Fig. S2**). Type II IFN (IFN-γ) was the most induced cytokine. Several other Th2 cytokines (IL-4/5/13) were also highly induced (**Fig. S2**). Notably, none of the Type I IFNs (IFN-α/β/ω) were induced by IIV (**Fig. S3**, blue bars, with the names of Type I IFNs marked).

**Fig. 4:**
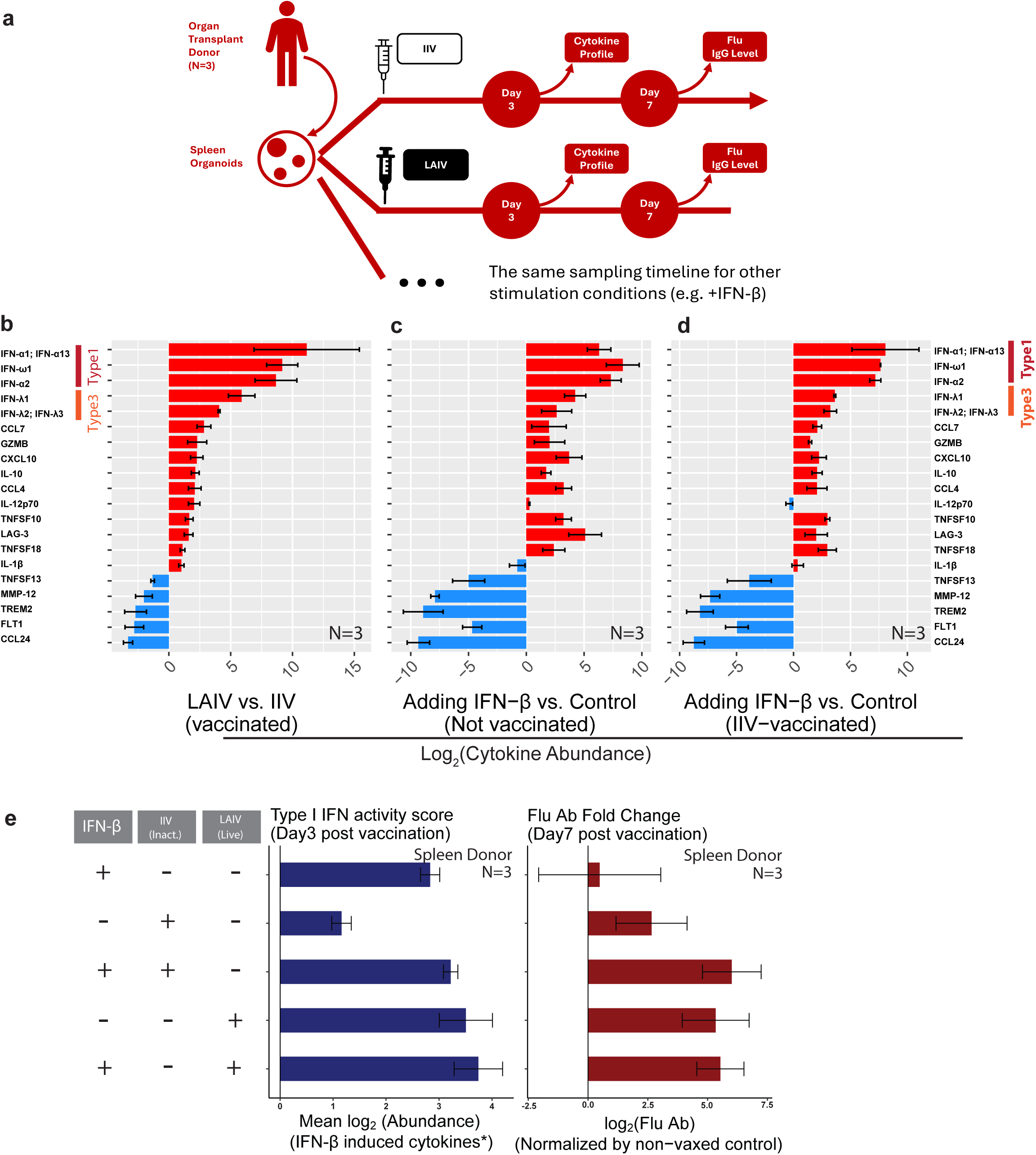
Type I IFNs drive an immune activation program distinguishing the response induced by live versus inactivated vaccines. **a)** We stimulated the spleen organoids derived from 3 transplant organ donors with IIV, LAIV, IFN-β, or combinations (see below). **b-d)** Cytokines differentially induced by LAIV versus IIV (FDR <0.05 and fold change >2) at Day 3 Post-VAX were plotted. Different stimulation conditions were noted in the bottom. **e)** We examined the Type I IFN activities **(left)** and flu antibody responses **(right)** of organoids under different stimulation conditions.

Next, we examined the cytokine-induction profile of another type of influenza vaccine, the live attenuated influenza vaccine (LAIV). Unlike the inactivated format, the live vaccine contains a mutated influenza virus capable of completing a few cell cycles in a human cell. Both IIV and LAIV are FDA-approved (but for different age groups and administration routes). The cytokine-induction profile of the live attenuated vaccine (LAIV) and inactivated vaccine (IIV) shared a strong induction of Th2 cytokines (such as IL-5, IL-13, IL-4), Th9 cytokine (IL-9), and Type II IFN (**Fig. S2-3**). However, the live attenuated vaccine also induced a unique and prominent Type I and III IFN response, with concentrations up to ∼1,000 times (10 log2) higher than IIV (**Fig. 4b and Fig. S3**). **S4**). Meanwhile, the live attenuated vaccine also induced a broad spectrum of other cytokines, including IL-10, IL-12, CCL-7, CXCL-10, etc. (**Fig. 4b**).

To examine the regulation of other cytokines by Type I IFN, we added IFN-β to the non-vaccinated spleen organoid controls. Strikingly, the cytokine profile induced by IFN-β almost overlapped with the differentially induced cytokines between the live and inactivated vaccine: 14 out of 15 LAIV-IIV differentially upregulated cytokines were also induced by IFN-β. In comparison, 4 out of 5 LAIV-IIV differentially downregulated cytokines were also suppressed by IFN-β (**Fig. 4c**). Similar cytokine-induction profile overlaps also persisted when we added IFN-β into the IIV-vaccinated controls and compared (**Fig. 4d**). As a result, the cytokine profile of the organoids treated with IFN-β phenocopied the live vaccine (**Fig. S4**).

When we added IFN-β to the spleen organoids, it induced the expression of other Type I IFNs (**IFN-**α**, IFN-**ω, **Fig. 4c-d**), which bind to the same receptor (IFNAR1/2) and induce the same signaling pathway. This observation suggests a positive feedback loop between the induction of Type I IFNs. These data also show that Type I IFN is a key regulator of the live-vaccine-specific cytokine profile, whose abundance differentiates the inactivated versus live vaccine responses (also including the antibody response, see below). Type I IFN triggers positive feedback on its expression and a cascade of other cytokines.

Next, we examined the functionality of Type I IFN (IFN-β) to adjuvant antibody responses. Due to the existence of multiple Type I IFNs, the concentration of a single protein could not represent the Type I IFN activity. We measured the global Type I IFN activity by taking the geometric mean of the abundance of all cytokines significantly induced by IFN-β. The Type I IFN activity scores showed that the inactivated vaccine was defective in inducing Type I IFN activity, which was rescued by IFN-β addition (**Fig. 4e**). While the live vaccine efficiently induced Type I IFN activity, adding more IFN-β did not further enhance it, suggesting saturation (**Fig. 4e**). Importantly, the antibody production in Day 7 post-VAX rose alongside the Type I IFN activity upon antigen challenge (**Fig. 4e**). Adding type I IFN (IFN-β) reshaped the downstream global cytokine profile to the live-vaccine-like state. Ultimately, adding IFN-β increased the antibody titer, making it resemble LAIV.

### IFN-**β** regulates the human vaccine response among older adults (age > 65y) receiving an insufficient dose of antigens

The association studies in human cohorts (**Fig. 1-2, and S5a**) and functional studies in tonsil and spleen organoids (**Fig. 4**) support that IFN-β is a natural adjuvant underlying flu vaccine response variability in humans. Interestingly, we found considerable heterogeneity in IFN-β’s correlation with post-VAX antibody response across the subpopulations (**Fig. S5a**). The heterogeneity was primarily due to age differences, with the correlation only reaching significance in older adults (>65 y). Among older adults, the correlation was more pronounced in those who received low-dose vaccines. When the low dose was found suboptimal for older adults ^44^ the cohort studies followed the most current knowledge at the time and switched the older adults to a version with high-dose antigen in the 2014-2015 season. Concurrent to this change, the association between baseline IFN-β concentration and antibody response disappeared (**Fig. S5b-c and Fig. S6**).

The age-dependent correlation between pre-VAX IFN-β abundance and post-VAX response prompted the hypothesis that certain aging-associated conditions may underlie this population’s IFN-β abundance and vaccine response variation. One of the cohorts in our analysis (the Stanford-Ellison Cohort, **Fig. 1b**) surveyed medical history annually. The study recorded 648 clinical events from 135 subjects, 628 of which occurred in the older adults (>65 y). Among them, nine types of clinical events accumulated more than 10 incidences in the cohort. We asked whether the serum IFN-β abundance changed within 2 years (before or after) a disease diagnosis. The most significant association identified is from osteoarthritis (**Fig. S7a**). Osteoarthritis is a highly prevalent condition in older adults, with 20 incidences in the cohort (compared to 3 cases of rheumatoid arthritis). While traditionally deemed a degenerative disease, a significant inflammatory^45^ (or infectious^46–48^) component has been identified in osteoarthritis. We leveraged the longitudinal nature of the study and examined the temporal correlation between serum IFN-β abundance and osteoarthritis incidence. The IFN-β abundance was significantly elevated from the population mean up to 4 years before diagnosis and lasted up to 5 years after (**Fig. S7b-d**). The temporal correlation was robust, retained significance after outliers were removed (>= 3 standard deviations from the population mean) (**Fig. S7c**). Within the time frame, we found that the antibody response to the influenza vaccine was significantly elevated in subjects with osteoarthritis (**Fig. S7e**). Thus, a highly prevalent aging-associated condition, osteoarthritis (including its pre-clinical stage), was one of the modifiers of the IFN-β abundance and vaccine response. This further reinforces the conclusion that IFN-β is a natural adjuvant in human beings and a source of variability in vaccine responses.

### IL-12 and IL-21 represent a Type I IFN-independent pathway in live-vaccine-specific response

In the immune organoid screen, we identified numerous cytokine adjuvants capable of boosting antibody response (**Fig. 3b**). Strikingly, most of the cytokine adjuvants were induced to higher levels by the live vaccine (**Fig. 4b**), compared to the inactivated vaccine. This highlights the important role of cytokines induced by the live vaccine. Four cytokine adjuvants are either Type I IFN (IFN-α, IFN-β, or IFN-ω) or induced by Type I IFN (IL-10) (**Fig. 4c,d**). However, two potent cytokine adjuvants (IL-12 and IL-21, **Fig. 3b**) are not induced by Type I IFN in human immune organoids (data for IL-12 shown in **Fig. 4c,d**, and IL-21 shown in **Fig. 5a,b**), and we identified them as live-vaccine-specific cytokines outside of the Type I IFN pathway. Prior studies have shown that IL-12 is the most potent human cytokine at inducing IL-21-expressing CD4+ T cells, which are responsible for B cell “help” {Crotty, 2019 #6;Schmitt, 2016 #5}. Meanwhile, the live attenuated influenza vaccine, but not the inactivated one, is known to activate the Tfh cells preferentially in human tonsil organoids^34,49^. Indeed, in spleen organoids, IL-12 can induce IL-21 (**Fig. 5a**). Thus, at least two distinct cytokine pathways (Type I IFN vs. IL-12/IL-21) regulated human vaccine responses (**Fig. 5c**).

**Fig. 5:**
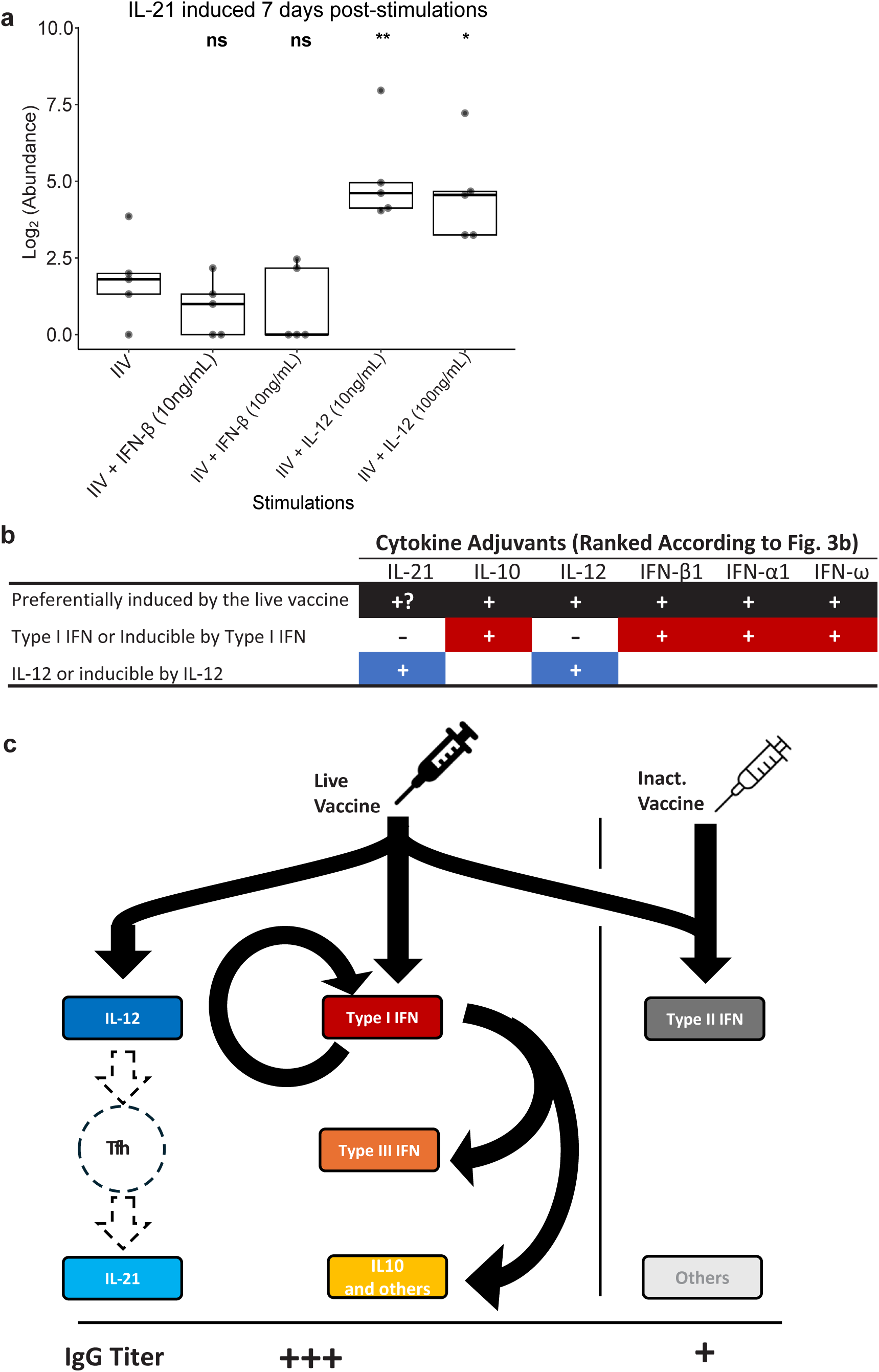
Two distinct cytokine pathways regulating vaccine responses in humans. **a)** IL-21 abundance measurement 7 days post-stimulation. **b)** The induction of cytokine adjuvants (capable of boosting vaccine responses, **Fig. 3b**). The induction profile is based on data in **Fig. 4c-d** and **Fig. 5a**. While the NULISA assays for the live vaccine in Figure 4 did not cover the IL-21 measurement, it is known that the live vaccine preferentially activates its main producer (Tfh) in human immune organoids. **c)**The model describes the different immune activation programs induced by the live and the inactivated influenza vaccines. We performed Wilcoxon ranking tests between different age groups. *, p<0.05; **, p<0.01.

### mRNA LNP can deliver cytokines to augment the quantity and breadth of antibody responses in mice

The above human studies showed that Type I IFN, IL-12, and downstream cytokines such as IL-21 could enhance antibody responses. Moreover, the screen quantitatively compared 19 cytokines, and we found IFN-β and IL-21 demonstrated the most potent adjuvant function at the lower concentration (1ng/ml, **Fig. 3b**). We next tested the functional relevance of these cytokine pathways in vivo using a murine model. Murine models complement the human model by offering a system where we can track the cross-tissue protein delivery (see below) and examine the antibody response at an organismal level. To this end, we employed a non-antigen-coding mRNA lipid nanoparticle (mRNA-LNP) platform in mice immunized with the inactivated influenza vaccine. The mRNA component of mRNA-LNPs is known to induce a strong Type I IFN response^51^, while its LNP component promotes the activation of Tfh cells^50^, the primary source of IL-21. This dual functionality enables mRNA-LNPs to mimic critical immunostimulatory features of a live vaccine, activating both Type I IFN and IL-21 pathways in vivo.

To evaluate this approach, we administered mRNA LNPs encoding GFP in combination with IIV in mice (**Fig. 6a**), which allowed easy tracking of the protein delivery (see below). Mice were boosted on Day 21 after the primary immunization, and serum samples were collected at multiple timepoints for antibody profiling. The IIV formulation included hemagglutinins (HAs) from human influenza strains (H1N1, H3N2, and B) circulating during the 2023-2024 season. Serum antibody responses were analyzed using a custom Luminex panel capable of detecting antibodies against both the in-vaccine (on-target) and out-of-vaccine (cross-reactive) hemagglutinins (HAs), including zoonotic strains such as H5N1 and H17N10 (**Fig. 6b**).

**Fig. 6:**
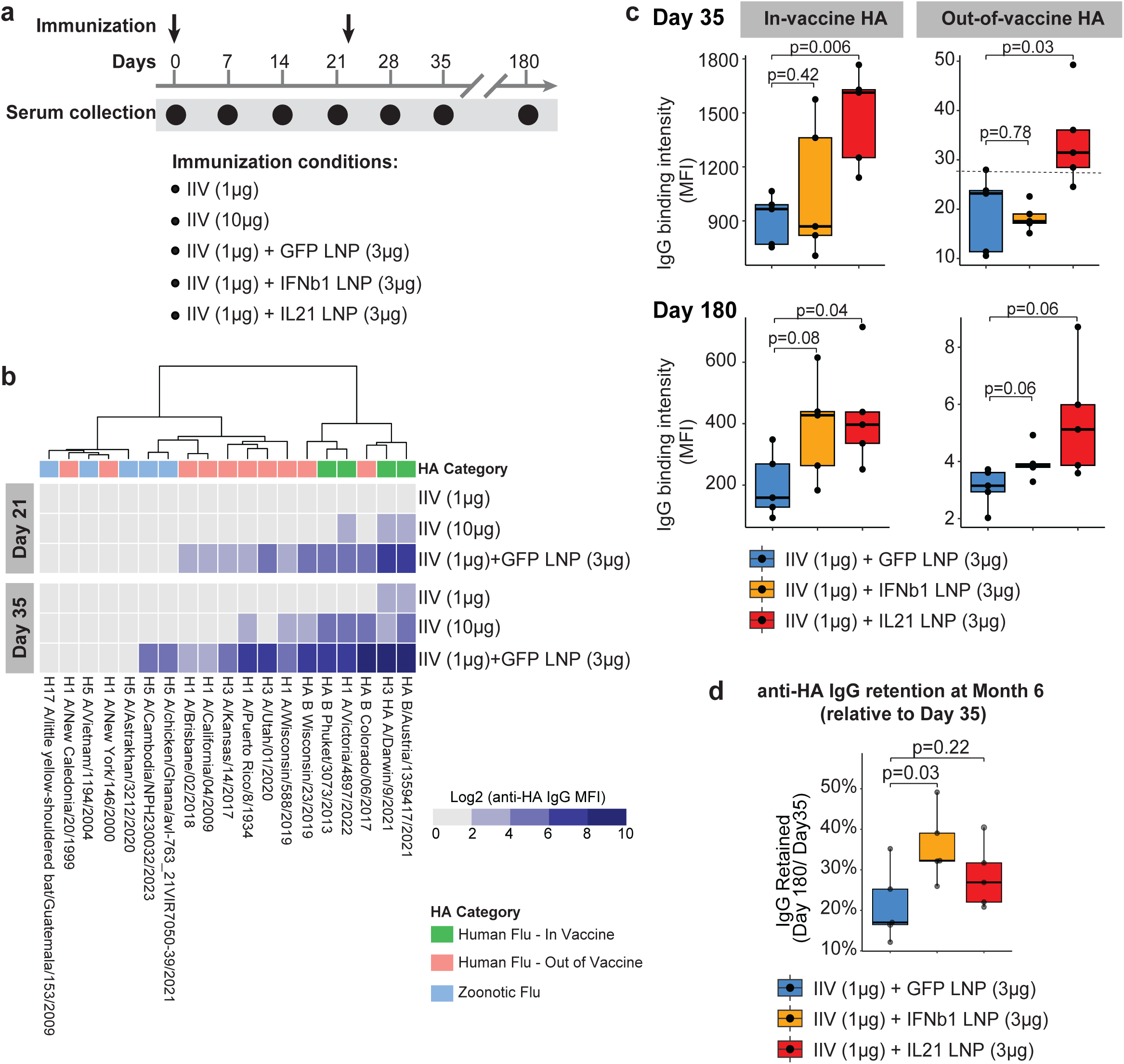
mRNA LNP augments the quantity and breadth of antibody response to the human inactivated influenza vaccine. **a)** Experiment scheme for testing the adjuvant functionality of mRNA lipid nanoparticles (mRNA-LNPs) encoding GFP or cytokines. Mice were primed on Day 0 and boosted on Day 21 as indicated. Sera were collected at the indicated timepoints for antibody titer detection. **b)** Heatmap showing serum IgG levels (as quantified by IgG binding intensities, Log2MFI) against a diverse panel of influenza hemagglutinins (HAs) from influenza, measured by custom Luminex bead assay on Day 21 and Day 35. HA antigens were categorized as Human Flu-In Vaccine, Human Flu-Out of Vaccine, or Zoonotic Flu as indicated by the top color bar. **c)** Comparison of anti-HA IgG titers across indicated groups at Day 35 and Day 180 post-vaccination, stratified by in-vaccine strains (left panel) and out-of-vaccine strains (right panel) (N=5 per group). The GFP mRNA-LNP group serves as the comparator. The horizontal dashed line indicates the IgG abundance against in-vaccine influenza strains induced by an unadjuvanted high-dose vaccine (10ug) at Day 35, whose antigen dose is at 10 times that of the adjuvanted condition. **d)** Percentage of anti-HA IgG retained at Month 6 relative to Day 35 (N=5 per group). In (c and d), the students’ t-test was performed with significance indicated. *, p<0.05; **, p<0.01.

At day 21 post-primary immunization, 1 μg IIV alone failed to elicit detectable antibody responses, while 10 μg IIV induced moderate titers against the vaccine, including in-vaccine HAs (B/Austria/1359417/2021, H3 A/Darwin/9/2021, and H1 A/Victoria/4897/2022, **Fig. 6b**). In contrast, 1 μg IIV combined with GFP mRNA-LNP markedly enhanced antibody titers against all in-vaccine HAs, and exceeded those elicited by 10 μg IIV alone. In addition, this group exhibited cross-reactive antibody responses to out-of-vaccine HAs (**Fig. 6b**). By day 35 (14 days post-booster), antibody titers increased further. The 1 μg IIV-alone group showed low but detectable two in-vaccine HAs B/Austria/1359417/2021 and A/Darwin/9/2021 (H3N2), while the 10 μg IIV group produced substantially higher titers to all in-vaccine strains, along with modest cross-reactivity. Notably, the group that received IIV plus GFP mRNA-LNP showed the highest antibody titers overall, including responses to Zoonotic HAs. The cross-reactive antibody titer against the two most recent H5N1 strains in this group was comparable to or higher than the antibody titer against in-vaccine strains induced by the high-dose non-adjuvanted influenza vaccines (10 μg, represented by the dashed lines) (**Fig. S8**). Thus, non-antigenic mRNA-LNPs boosted the quantity and the breadth of humoral responses to inactivated vaccines.

To examine the tissue localization of mRNA-LNP activity, we tracked GFP tissue distribution 24 hours post-injection. Fluorescence was detected at both the injection sites, muscles, and draining lymph nodes, with substantially higher intensity in the lymph nodes (**Fig. S9**). This suggested efficient delivery of mRNA and protein expression in secondary lymphoid tissues. This supports the possibility that cytokines encoded by mRNA-LNPs could act in lymph nodes locally to modulate vaccine response as an adjuvant. Thus, we next tested whether mRNA-LNPs encoding specific cytokines could enhance vaccine responses beyond the innate adjuvanticity of LNPs themselves (**Fig. 6a**). Mice were immunized with 1 μg IIV in combination with mRNA-LNPs encoding either mouse IFN-β or IL-21. At Day 35, inclusion of IL-21 mRNA LNPs significantly outperformed GFP mRNA-LNPs in boosting antibody titers against both in-vaccine and out-of-vaccine HA antigens (**Fig. 6c**). At the same time, IFN-β mRNA-LNPs showed no advantage at this time point. However, at six months post-immunization, both IL-21 and IFN-β mRNA LNP resulted in higher antibody titers than the GFP mRNA-LNPs (**Fig. 6c**). Comparison of titers between the 6-month and Day 35 revealed that IFN-β mRNA-LNPs slowed the waning of antibody response and provided greater durability than IL-21 (**Fig. 6d**).

These results demonstrate that cytokines IFN-β and IL-21 can serve as effective vaccine adjuvants to enhance antibody responses. Using mRNA-LNPs to deliver these cytokines in vivo provides a flexible strategy to boost the immunogenicity of inactivated vaccines. This approach offers a rational path to improve the efficacy of inactivated or subunit vaccines, particularly those with limited innate immunogenicity.

## Discussion

This study identified several cytokines (IL-18, GROα/CXCL1, IFN-β) whose pre-vaccination concentration correlated with the specific antibody response to influenza vaccines. Identifying baseline predictors of an individual’s vaccine response is a key goal for systems vaccinology^7–9^. Prior studies have investigated the correlation between transcriptome signature and influenza vaccine response^10,51,52^. Cytokines are likely molecular drivers for many observed transcriptome signatures. In this study, we combined human cohort studies with *in vitro* analyses using immune organoids to identify cytokines that augmented flu vaccine responses. We stratified subjects by their pre-VAX HAI flu antibody titer and focused only on those with low pre-VAX HAI flu antibody titer. This design was based on our finding that the biological signals (such as aging effects) were concentrated in the population with low pre-VAX HAI flu antibody levels (**Fig. 1c**). This finding is in line with prior work that showed subjects with low pre-VAX HAI flu antibody launched more robust interferon responses after the vaccination, which may be attributed to less vaccine antigen clearance and thus more vaccine “take”. We demonstrated that stratification is essential to detect the correlation between pre-VAX cytokine abundance and post-VAX antibody response. The stratification design may be applied to other vaccine research.

Combining correlative and interventional studies, we found that IFN-βis a natural adjuvant capable of boosting antibody response in the human population (**Fig. 1,2, and 5**). The IFN-related gene expression signature (induced by either Type I or II IFN^53^) at pre- or post-vaccination time points correlated with Day 28 antibody responses^10,11,13,14^. A previous study reported a shared transcriptome signature between vaccine high-responders in patients with systemic lupus erythematosus, a disease characterized by chronic activation of Type-I IFN signaling^11^. Meanwhile, higher Type II IFN production (by CD8^+^ T cells) has been reported to correlate with influenza vaccine responses in COVID-19 recoverees^12^. Also, SLE patients who did not respond to the SARS-Cov2 vaccine had almost no interferon-γ responses following a booster^54^. The relative contribution of Type I or Type II IFNs to vaccine responses has not been tested in a human immune model. We performed a functional screen of cytokine adjuvants in immune organoids. We found that Type I IFNs, but neither Type II IFN nor IL-18 (a potent inducer of Type II IFN^55^), could enhance influenza vaccine responses. Whether the finding is specific to the influenza vaccine or can be generalized to other vaccines is worth further investigation. Our findings suggest that Type I IFN is a master regulator specific for the live vaccine response, upstream of most other cytokine activation programs. Live vaccines (e.g., measles vaccine, LAIV) tend to generate a more robust human protection than their inactivated counterparts^56–59^. Comparative studies in mouse models using influenza or rabies vaccines (live versus inactivated) also confirmed the finding^60,61^. Extended antigen exposure can augment antibody responses^62^. Additionally, the live vaccine may activate a specific immune program to drive an augmented antibody response. In spleen organoids, we performed a comparative study of 250 cytokines using a high-throughput technology with DNA-barcoded antibodies (NULISA). The assay identified 15 live-vaccine-specific cytokines (**Fig. 4b**). Adding a single cytokine, type I IFN (IFN-β), recapitulates most of the live-vaccine-specific immune activation program (**Fig. 4c-d)**, including other type I IFNs and another cytokine adjuvant, IL-10 (but not IL-12 or IL-21, see below). Moreover, adding Type I IFN boosted the inactivated vaccine’s antibody induction quantity to a level similar to the live vaccine (**Fig. 4e**). Consistent with these findings in humans, mice lacking IFNAR1 or STAT1 exhibit impaired viral clearance, altered isotype class switching, and reduced IgG2c and IgA production after influenza vaccination^63–65^. We also examined the underlying heterogeneity within the large cohorts. Baseline IFN-β abundance correlated with antibody response primarily in old adults (>65y) receiving low-dose antigen (**Fig. S5a**). Moreover, in older adults, osteoarthritis incidence is associated with a higher IFN-β baseline and antibody response to the inactivated vaccine (**Fig. S7**). We noted that medical history recorded in the cohort relies on self-report during the annual visit, which is a limitation of the study. Also, due to the sample size limitation, we could not examine the correlation between vaccine response and other well-established Type I interferonopathies with a lower incidence rate in the population (such as lupus).

Our systemic screen also identified a second pathway (IL-12/IL-21) that is Type I IFN-independent (**Fig. 5b-c**). Human type I interferon deficiency-due to inborn errors (e.g., IFNAR1/2, IRF7 mutations) or neutralizing auto-antibodies-impairs antiviral immunity and underlies severe infectious diseases (influenza^66^, COVID-19^67^, herpes simplex^68^) or severe adverse reactions to live vaccines (measles and yellow fever^69^). Despite profound defects in the innate antiviral defense, individuals with type I IFN deficiency can still mount humoral responses. For example, YFV-neutralizing antibodies in IFNAR1-deficient patients can develop following vaccination and exposure^70^, indicating that B cell priming and antibody production can proceed independently of intact type I IFN signaling. We found that in human spleen organoids IL-12 and IL-21 were outside the Type I IFN pathway (not induced by adding Type I IFN, **Fig. 4c,d and Fig. 5**). We also demonstrated that IL-21 can be induced by IL-12 (**Fig. 5**). Meanwhile, in a recent report, IL-12 can also regulate B cell function directly^71^. Thus, the live vaccine augments a robust antibody response by activating diverse cytokines (at least six different cytokine adjuvants, **Fig. 5b**) in orthogonal pathways (Type I IFN vs. IL-12/IL-21). In addition, IL-12 and IL-21 were outside the Type I IFN pathway (not induced by adding Type I IFN, **Fig. 4c,d and Fig. 5a,b**). We also demonstrated that IL-21 can be induced by IL-12 (**Fig. 5a**), potentially due to Tfh activation^21^. Meanwhile, in a recent report, IL-12 can also regulate B cell function directly^71^. Thus, the live vaccine augments antibody responses by activating diverse cytokines (at least six different cytokine adjuvants, **Fig. 6b**) in orthogonal pathways (Type I IFN vs. IL-12/IL-21). RNA replication of the influenza virus (occurred for a few cycles for the live attenuated vaccine) results in 5′-triphosphate ssRNA, which can activate pattern recognition receptors (RIG-I)^72^. Further investigation of the innate immunity sensing process can reveal the upstream mechanisms for the differential cytokine activation program between the live and inactivated vaccines. Meanwhile, the interaction of the orthogonal cytokine pathways (Type I IFN vs. IL-12/IL-21) on the quantity and quality (breadth) of antibody responses warrants further investigation.

In line with our findings in human cohorts and organoids, we found that non-antigen-coding mRNA LNP, a potent inducer of both Type I IFN and IL-21, substantially enhanced the quantity and breadth of the antibody response to the inactivated influenza vaccine. Non-antigen coding mRNA LNP was so potent that the H5N1 HA antibody levels against the two most recent H5N1 strains induced by the adjuvanted vaccine (1µg IIV + 3µg GFP mRNA LNP, Fig. 6c) were comparable to the human influenza HA antibody levels induced by the high-dose non-adjuvanted vaccine. Our results aligned with the prior finding that lipid nanoparticles exert a potent adjuvant effect by augmenting Tfh expansion^50^. Importantly, including IL-21 mRNA or IFN-β further augmented the antibody response already potently adjuvanted by mRNA-LNP. While mRNA-LNP has been widely used to encode antigens (such as in COVID-19 vaccines), its potential for delivering cytokine adjuvants has not been fully explored. Beyond IL-21, a recent study reported that mRNA LNP encoding IL-12 enhanced the vaccine response to COVID-19 antigens^73^. Both IL-21 and IL-12 showed a potent adjuvant effect in our organoid screen of 19 cytokines (**Fig. 3b**). However, the lipid components of mRNA-LNP vaccines, particularly the ionizable and PEGylated lipids, can induce potent and broad inflammatory effects, including the activation of complement, and the release of multiple cytokines, including IL⍰1α, IL⍰1β, IL⍰6, IL⍰8, TNF⍰α, and IFN⍰γ^74–76^. Many of these cytokines have been linked to febrile or other adverse effects^77,78^. Future work can explore the possibility of cytokine mRNA adjuvants encapsulated with immunologically inert lipids to improve precision in adjuvant engineering.

Our study focused on antibody responses. Most cohorts included only assayed antibody quantity/titers, which correlated with protection for influenza infections^10,51,58^. Using a Luminex platform capable of detecting both in-vaccine and cross-reactive antibodies, we also measured the impact of mRNA-LNP and cytokines on breadth of antibody response (**Fig. 6**). Future studies can investigate the cytokine regulation of cellular immunity in human vaccine responses. Overall, our study identified a cytokine, IFN-β that acts as a natural adjuvant which contributes significantly to vaccine response variability in human populations with low baseline Ab titers and is also a key regulator of the live-vaccine-specific immune activation program. Moreover, we used a functional systems immunology approach to screen the adjuvanticity of numerous cytokines in immune organoids. The screen identified six cytokine adjuvants involved in the live-vaccine-specific antibody response, in at least two different pathways. This knowledge can inform the design of adjuvants that mimic the live-vaccine-specific effect without the risk of viral replication.

## Acknowledgements

Mark Davis is supported by NIH (CCHI U19 2U19AI057229-16) and Open Philanthropy. Bill Chen and Ashley Smith are supported by the Medical College of Wisconsin Start-up Fund, the Women’s Health Research Program, and the Sharon L. La Macchia Innovation Fund. Bill Chen was also supported by the Life Science Research Foundation (sponsored by Eli Lilly). John Heath was supported by the National Science Foundation Graduate Research Fellowship Program (NSF GRFP). Paul J. Utz was supported by the Henry Gustav Floren Trust, the Stanford Department of Medicine Team Science Program, the Stanford Medicine Office of the Dean, and the National Institutes of Health (NIH R01 AI182319-02). Ying Lu was supported by the NIH 1UL1TR003142. Holden Maecker was supported by the CCHI (NIH grant # 2U19AI057229) and HIPC (NIH grant # 5U19AI090019). The authors would also like to acknowledge the assistance of the Stanford Microfluidics Foundry.

