## supplementary figures for "Cytokine Regulation of Human Antibody Responses to Influenza Vaccines"

### Figure S1

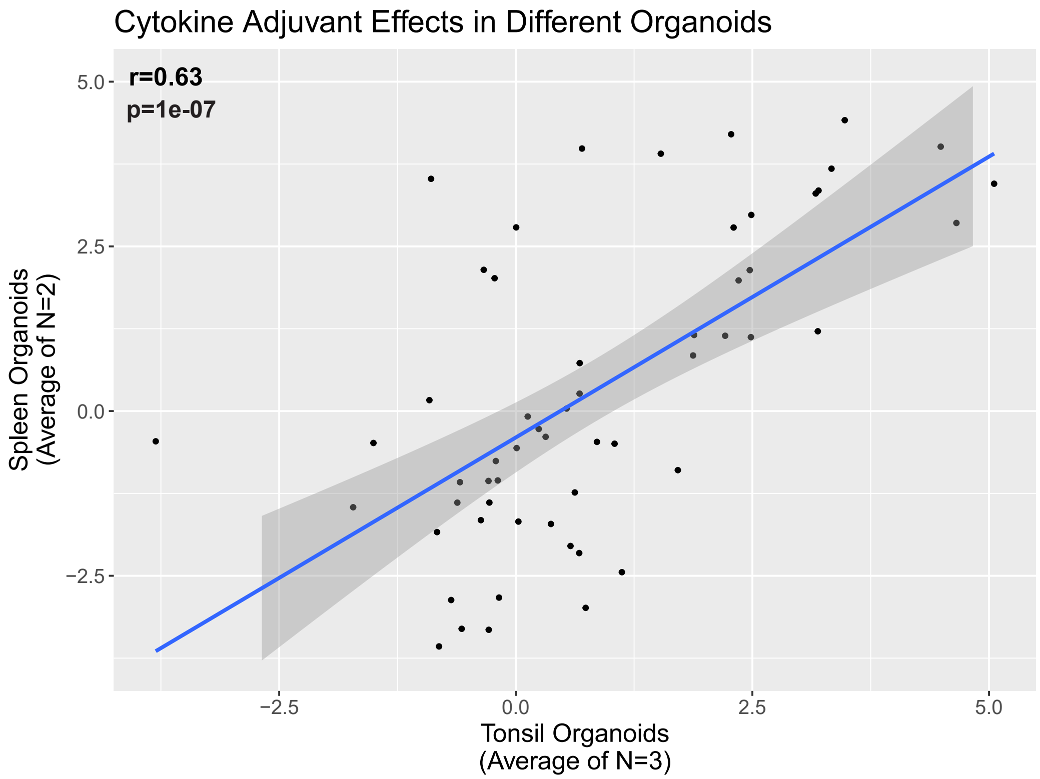


**Fig. S1: Cytokines demonstrated similar IIV-adjuvating effects in spleen and tonsil organoids.**

This figure is related to **Fig. 3**. We harvested 3 tonsils and 2 spleens from donors and vaccinate the organoid cultures with IIV. We added 19 different cytokines together with the vaccine (IIV), and for each cytokine, we tested 3 different concentrations ranging from 1ng/ml to 100ng/ml. We measured the antibody production (relative to non-cytokine-added IIV only controls) in Day 7. The adjuvant effects of 19 cytokines at 3 different concentrations (total 57 points) are plotted on the scatter plot, with the x axis representing the averaged value for tonsil organoids and y axis for spleen organoids. The Pearson's correlation coefficient and p values are listed.

### Figure S2

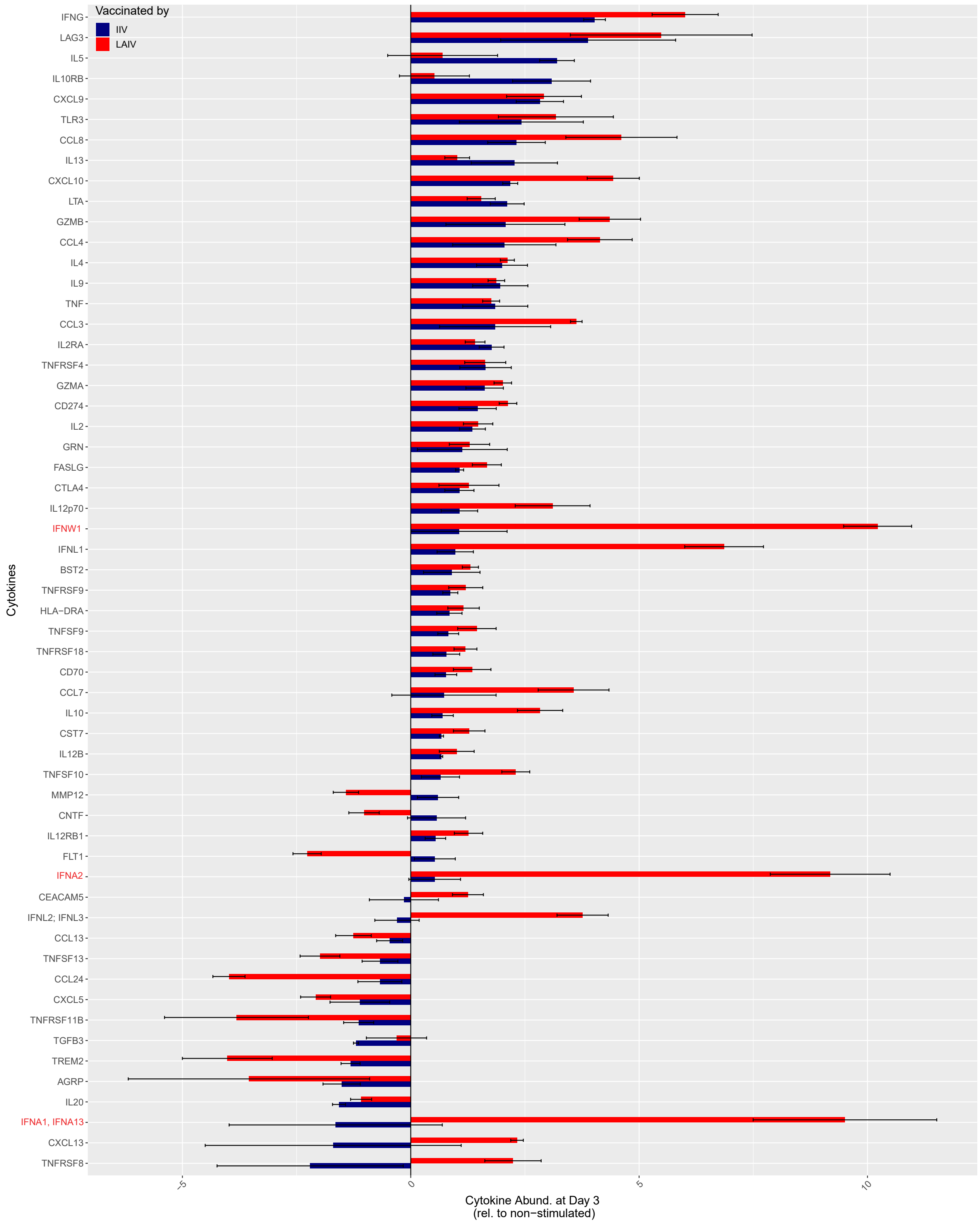


**Fig. S2: The cytokine profiles induced by IIV or LAIV in spleen organoids**

This figure is related to **Fig. 4**. We measure the cytokine profiles using supernatants of the spleen organoids (N=3) culture at Day 3 post-VAX. All cytokines significantly altered after vaccination by either vaccine (as compared with the non-stimulated) are shown. The Type I interferons (IFNs) are highlighted by red texts.

### Figure S3

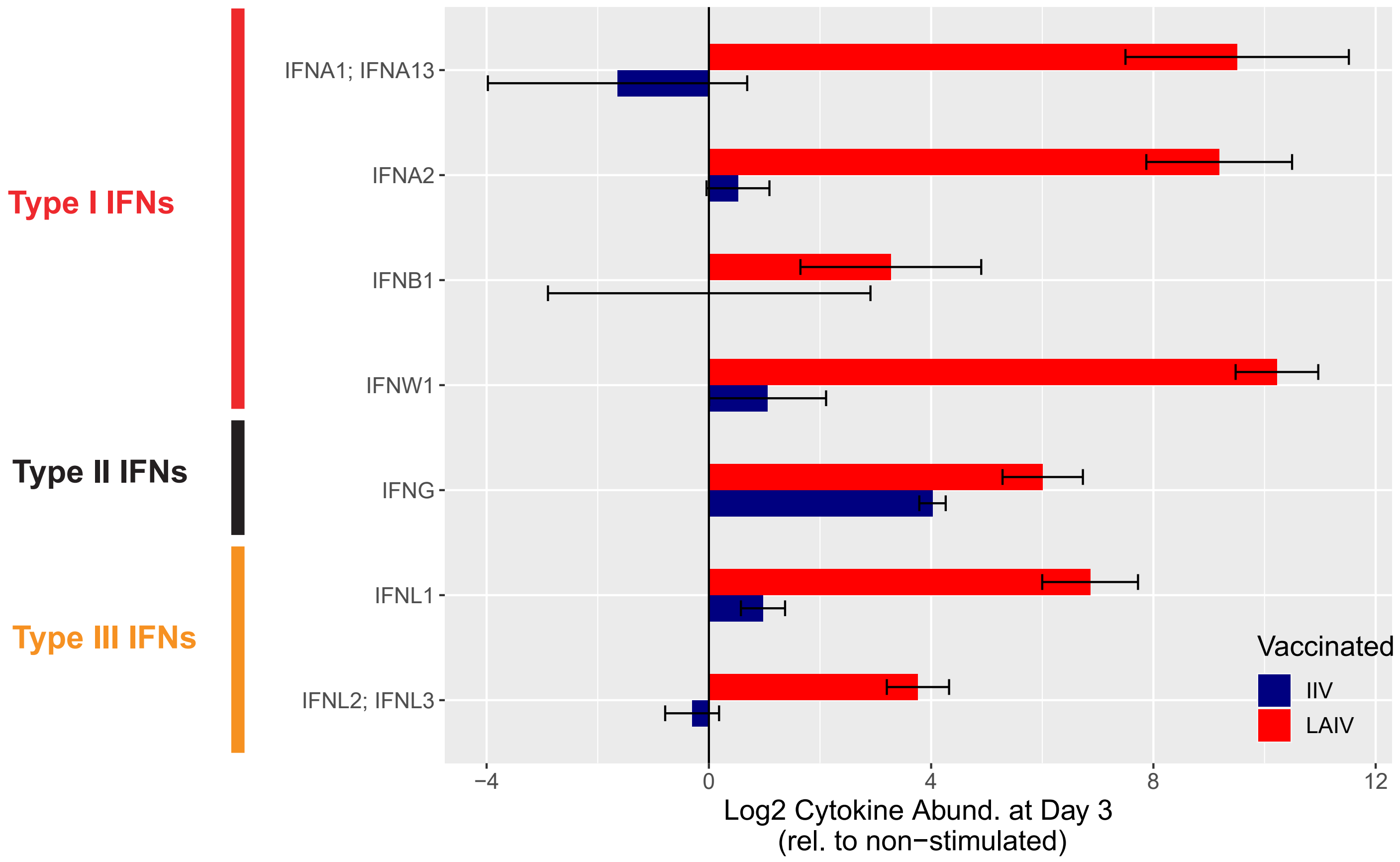


**Fig. S3: IFNs induced by IIV or LAIV in spleen organoids**

The figure is related to **Fig. 4**. The abundances of Type I/ II/ III cytokines in the supernatants of spleen organoids stimulated by either IIV or LAIV were reported.

### Figure S4

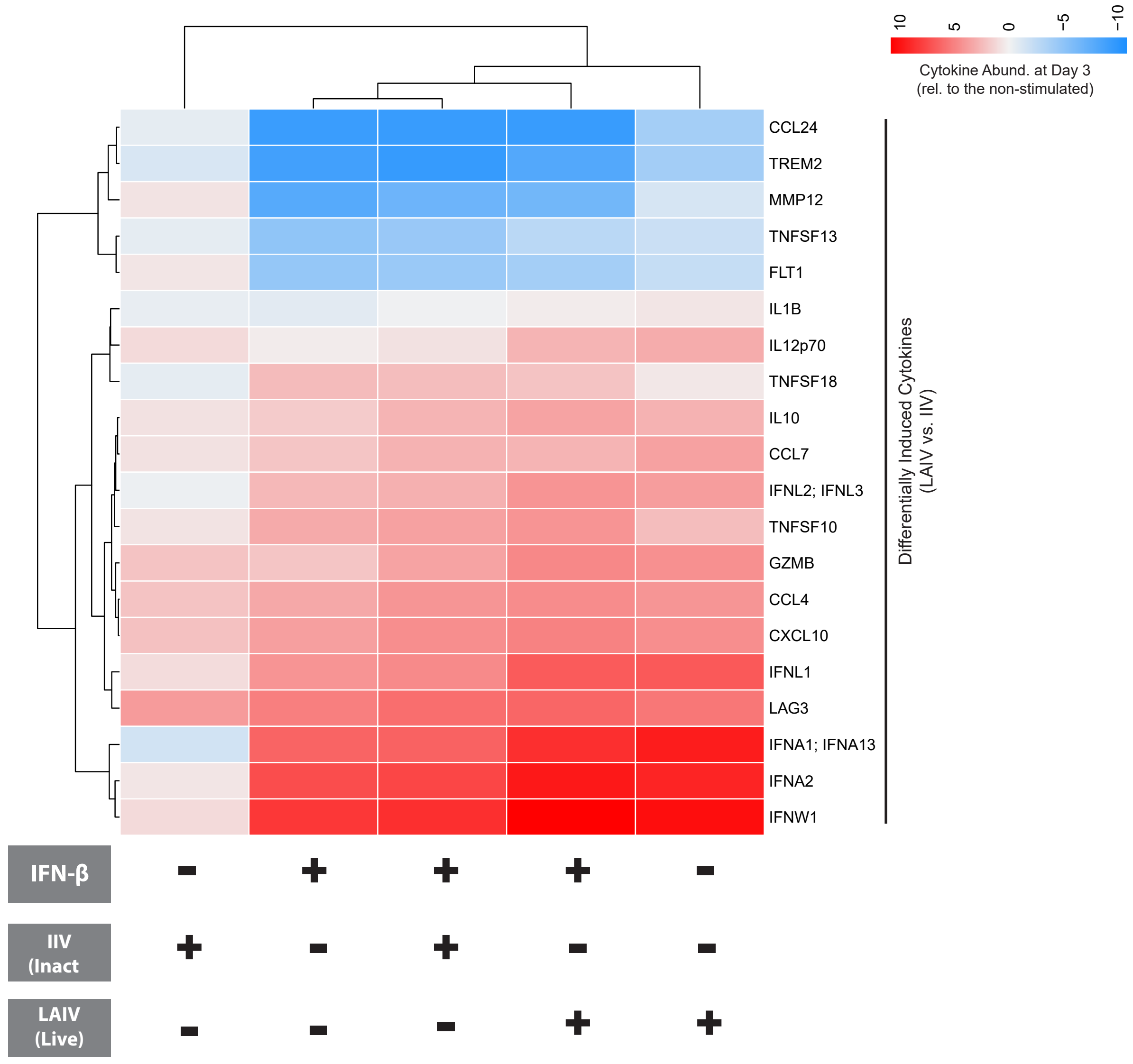


**Fig. S4: The cytokine profiles of spleen organoids under different stimulation conditions.**

This figure is related to **Fig. 4**. We measure the cytokine profiles using supernatants of the spleen organoids (N=3) culture at Day 3 post-VAX. Cytokines differentially induced by LAIV versus IIV (FDR <0.05 and fold change >2) are shown.

### Figure S5

**a**

**Correlation between Post-Vax Ab Resp. and Pre-Vax IFN-β Serum Abund.**
**Across Sub-populations**

| | Age (y) | Vax(IIV) Dose | Gender | Pre-VAX Ab Quantile | Trial/Cohort | Flu Season | Subject(N) |
|---|---|---|---|---|---|---|---|
| | [ 2,18) | 15 ug | Male | 25-50% | Flu Vaccine Study No.18 | 2010 | 6 |
| | [ 2,18) | 15 ug | Male | 25-50% | Flu Vaccine Study No.18 | 2013 | 9 |
| | [18,65) | 15 ug | Female | 0-25% | Flu Vaccine Study No.18 | 2010 | 11 |
| | [18,65) | 15 ug | Female | 25-50% | Flu Vaccine Study No.15 | 2009 | 6 |
| | [18,65) | 15 ug | Female | 25-50% | Flu Vaccine Study No.15 | 2015 | 5 |
| | [18,65) | 15 ug | Female | 25-50% | Flu Vaccine Study No.17 | 2009 | 6 |
| | [18,65) | 15 ug | Female | 25-50% | Flu Vaccine Study No.17 | 2010 | 8 |
| | [18,65) | 15 ug | Female | 25-50% | Flu Vaccine Study No.18 | 2010 | 16 |
| | [18,65) | 15 ug | Female | 25-50% | Flu Vaccine Study No.28 | 2014 | 7 |
| | [18,65) | 15 ug | Male | 0-25% | Flu Vaccine Study No.15 | 2015 | 5 |
| | [18,65) | 15 ug | Male | 0-25% | Flu Vaccine Study No.18 | 2010 | 6 |
| | [18,65) | 15 ug | Male | 25-50% | Flu Vaccine Study No.15 | 2009 | 9 |
| | [18,65) | 15 ug | Male | 25-50% | Flu Vaccine Study No.18 | 2010 | 9 |
| | [65,99] | 60 ug | Female | 0-25% | Flu Vaccine Study No.15 | 2015 | 11 |
| | [65,99] | 60 ug | Female | 25-50% | Flu Vaccine Study No.15 | 2015 | 7 |
| | [65,99] | 60 ug | Female | 25-50% | Flu Vaccine Study No.17 | 2010 | 6 |
| | [65,99] | 60 ug | Male | 0-25% | Flu Vaccine Study No.15 | 2015 | 6 |
| | [65,99] | 60 ug | Male | 25-50% | Flu Vaccine Study No.15 | 2014 | 9 |
| | [65,99] | 60 ug | Male | 25-50% | Flu Vaccine Study No.15 | 2015 | 5 |
| A | [65,99] | 15 ug | Female | 0-25% | Flu Vaccine Study No.15 | 2009 | 6 |
| B | [65,99] | 15 ug | Female | 0-25% | Flu Vaccine Study No.18 | 2009 | 9 |
| C | [65,99] | 15 ug | Female | 25-50% | Flu Vaccine Study No.15 | 2009 | 14 |
| D | [65,99] | 15 ug | Female | 25-50% | Flu Vaccine Study No.15 | 2010 | 12 |
| E | [65,99] | 15 ug | Female | 25-50% | Flu Vaccine Study No.15 | 2013 | 10 |
| F | [65,99] | 15 ug | Female | 25-50% | Flu Vaccine Study No.17 | 2009 | 5 |
| G | [65,99] | 15 ug | Female | 25-50% | Flu Vaccine Study No.18 | 2009 | 7 |
| H | [65,99] | 15 ug | Male | 25-50% | Flu Vaccine Study No.15 | 2009 | 10 |
| I | [65,99] | 15 ug | Male | 25-50% | Flu Vaccine Study No.15 | 2013 | 8 |

| Summary | Sample (N) |
|---|---|
| | 228 |

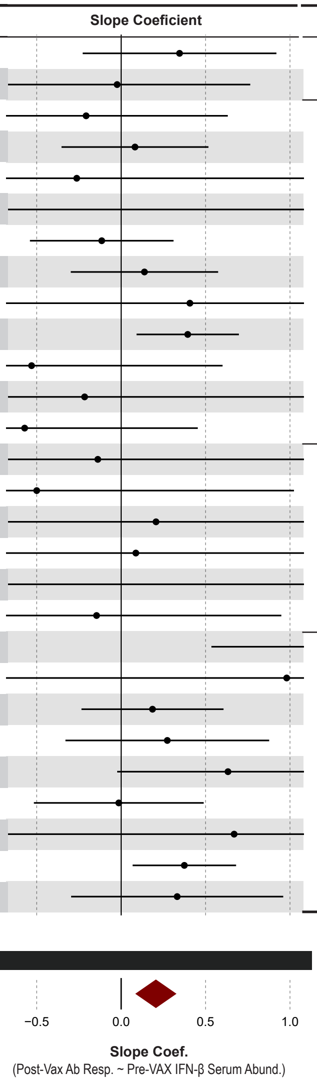


**b**

**Statistics by Group**

| Group | Summary Significance |
|---|---|
| Children | n.s. p= 0.36 |
| Young Adults | n.s. p= 0.21 |
| Old, High Dose | n.s. p= 0.92 |
| Old, Low Dose | ** p= 0.0011 |
| All | *** p= 0.00096 |

**c**

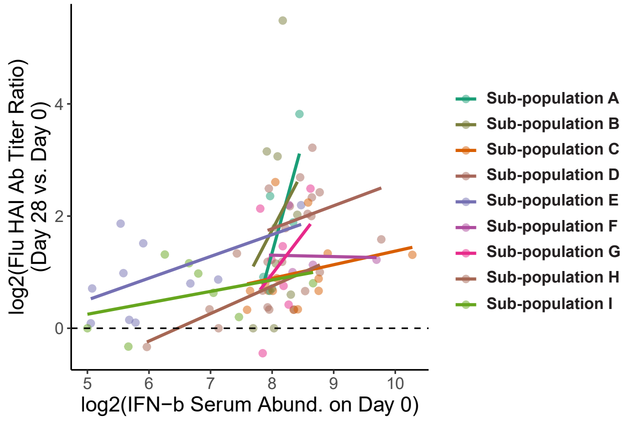


**Fig. S5: Pre-VAX IFN-β serum abundances correlates with Post-VAX antibody responses in the elderly receiving low-dose flu vaccines.**

We divided the dataset by confounders (age, vaccine dose, gender, day 0 flu antibody quantile, cohort, flu season) into 28 sub-populations and performed correlations with each sub-population. **a)** The left panel described the characteristics of each one of the 28 sub-populations. The slope coefficients panel described the correlation within each sub-population, with summary statistics provided in the bottom. **b)** We summarized the 28 sub-population into 4 groups by age and vaccine doses. For each group, summary statistics are provided. Details about the correlation and meta-analysis can be found in supplementary methods. **c)** The correlation between IFN-β serum abundance and antibody response in the old receiving a low dose IIV. The sub-populations are marked accordingly in **a**. The IFN-β serum abundance data are not normalized here, with the batch effects visible. Also see **Fig. S6**.

### Figure S6

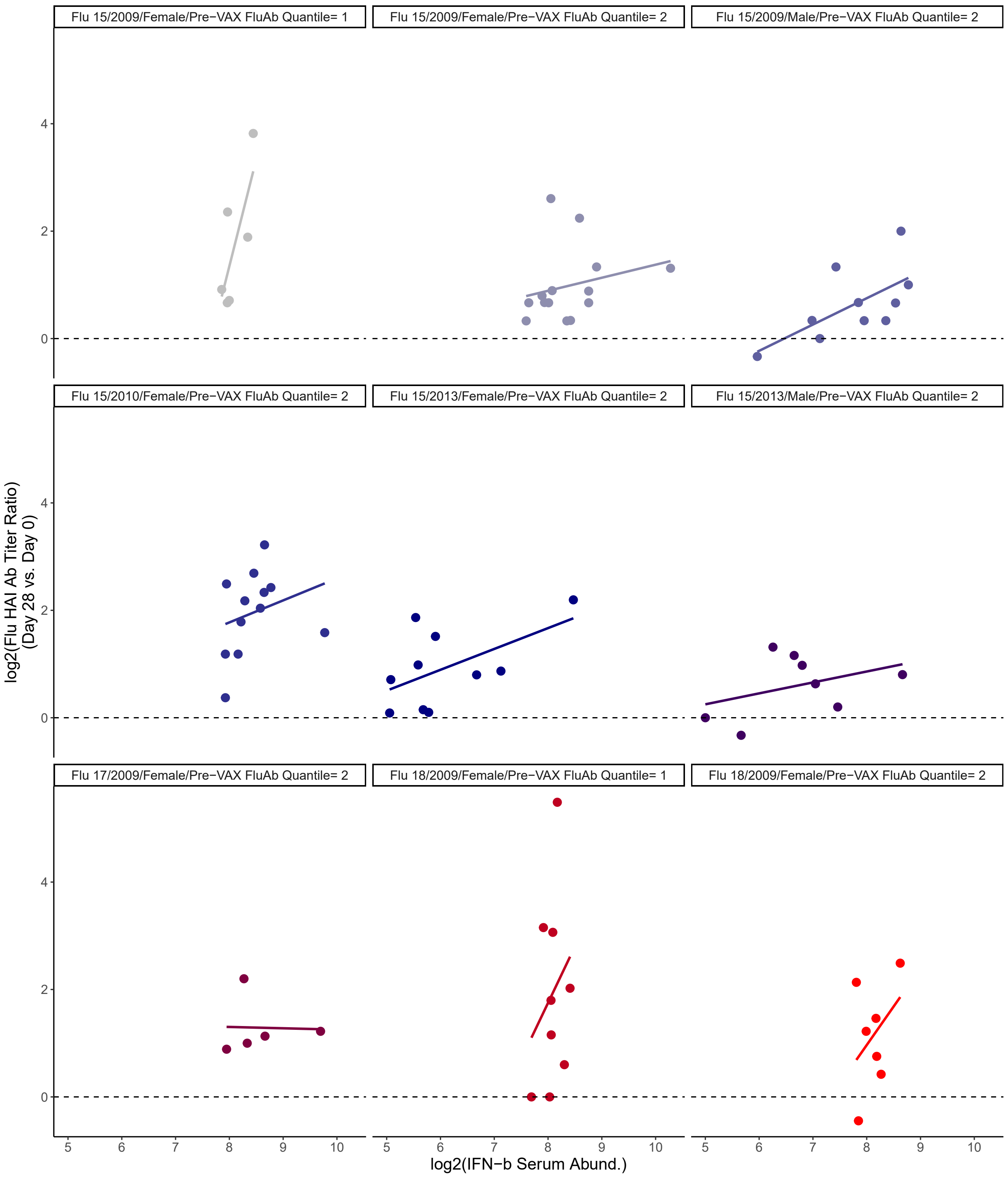


**Fig. S6: The correlation between pre-VAX IFNβ serum abundance and post-VAX flu antibody responses in the elderly receiving a low dose flu vaccine.**

This figure is related to **Fig. S5c.** The correlations (trends highlighted by the lines) of the 9 sub-populations in the “Old, low dose” group (**Fig. S5b, lower**) are plotted. The title of each panel corresponds to a sub-population listed in **Fig. S5a**.

### Figure S7

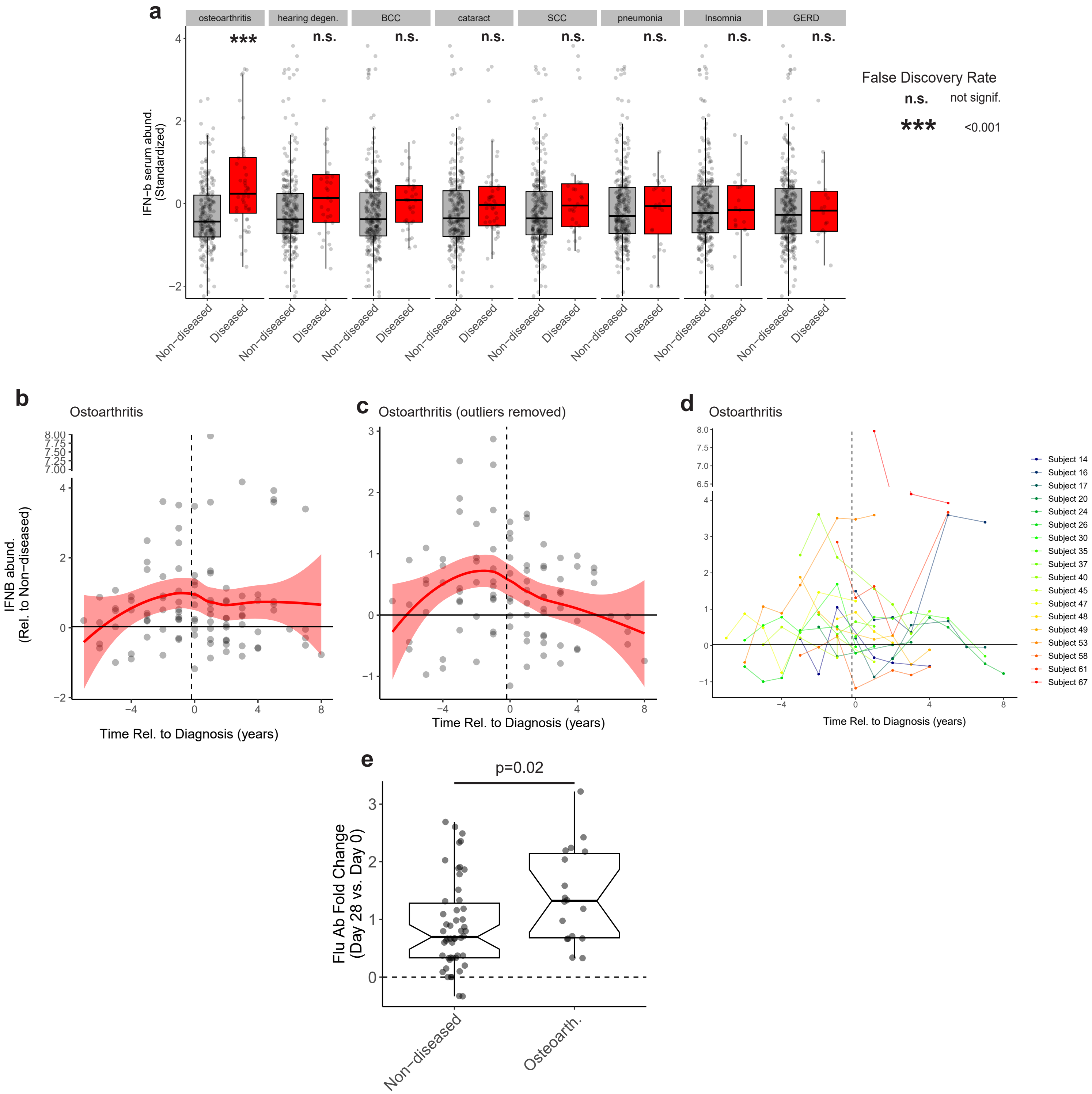


**Fig. S7: Osteoarthritis incidence modifies the IFN-β abundance and vaccine response**

The figure is related to **Fig. S5**. We examined the correlation between disease incidence, IFNβ serum abundance and flu vaccine antibody response in a longitudinal cohort (Flu Study 15, one of the 5 cohorts in **Fig. 1a**). This cohort collected medical histories annually. The analysis was conducted within the elderly (>65y). **a)** The IFNβ serum abundance between the diseased and non-diseased samples. The diseased samples are samples collected within (+/-) 2 years relative to a diagnosis. The non-diseased samples are from subjects without a medical history of the disease. The comparisons were performed by Wilcoxon ranking tests. **b-d)** Temporal profile of IFNβ serum abundance relative to the diagnosis of osteoarthritis within the cohorts. The loess smooth line is provided with confidence intervals. **c)** The outliers (standardized abundance >=3) were removed, and the temporal correlation retains. d) the temporal profile of individual patients diagnosed with osteoarthritis during the study. **e)** The flu HAI antibody responses (Day 28 vs. Day 0 ratio) in non-diseased and osteoarthritis subjects. Similar to Figs 2 and 3, the analysis was performed in subjects with a low level of pre-VAX flu antibodies (**Fig. 1c**). Refer to the supplementary methods.

### Figure S8

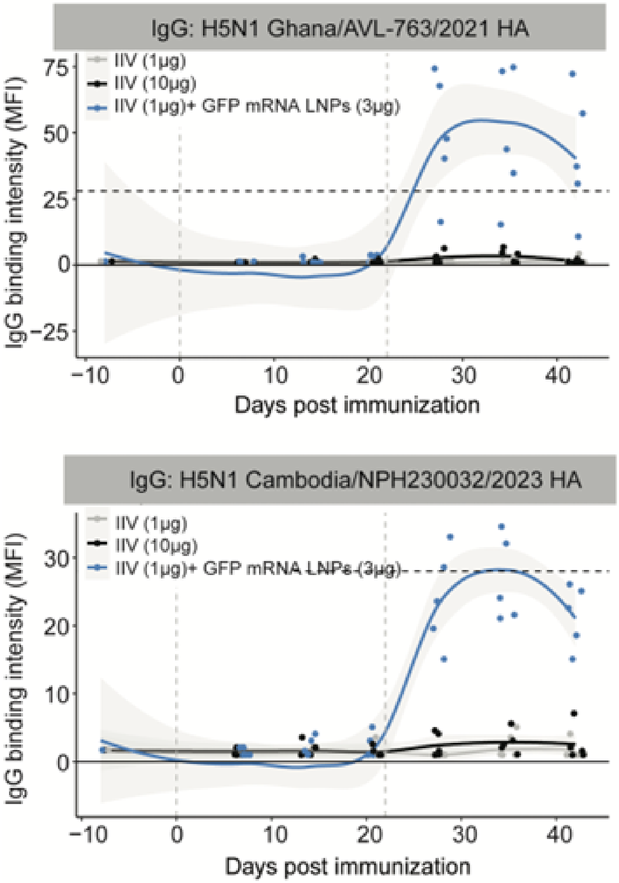


**Fig. S8: The kinetics of IgG responses to H5N1 HA antigens following IIV immunization with or without mRNA-LNP adjuvant.**

Serum IgG binding intensities (MFI) against recombinant H5N1 HA proteins from A/chicken/Ghana/AVL-763/2021 (top) and A/Cambodia/NPH230032/2023 (bottom) were quantified over time using a custom Luminex assay. Mice were immunized as shown in Figure 6a. The two vertical dashed lines indicate the timing of the prime and boost immunizations. The horizontal dashed line represents the IgG response (MFI) against in-vaccine influenza strains induced by unadjuvanted high-dose vaccine (10ug) at Day 35, whose antigen dose is 10 times that of the adjuvated condition..

### Figure S9

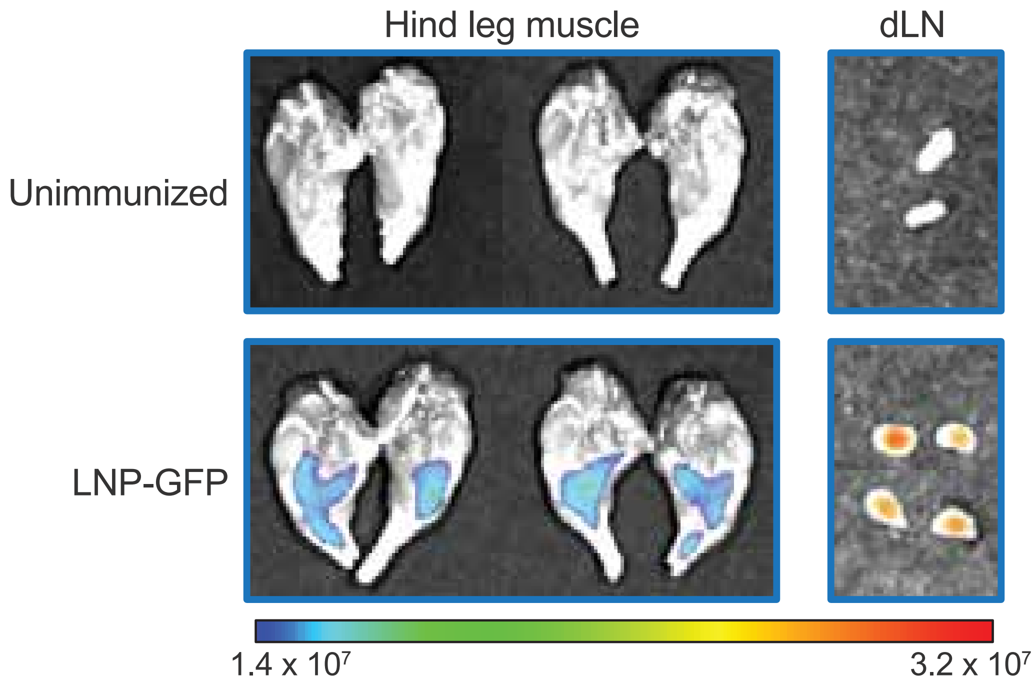


**Fig. S9: Fluorescence imaging of GFP in hind leg muscles and inguinal draining lymph nodes (dLNs) from unimmunized and LNP-GFP immunized mice.**

The images were taken 24 hours after injection. Data are representative of two independent experiments.
