## supplementary information for "Cytokine Regulation of Human Antibody Responses to Influenza Vaccines"

### Methods

#### Cohort Design and Data Retrieval

#### We retrieved human serum cytokine and antibody data from previous influenza vaccine studies conducted by the Stanford Human Immune Monitoring Center (HIMC). Only individuals who received non-adjuvanted, inactivated influenza vaccines (Fluzone®, standard or high dose) were included; subjects who received FluMist® (live attenuated) or Fluad® (MF59-adjuvanted) vaccines were excluded. Various laboratories measured hemagglutination inhibition (HAI) antibody titers between 2007 and 2015. To ensure data quality across years, we analyzed a longitudinal cohort (Flu Vaccine Study No. 15) to assess year-to-year consistency. The 2015 measurements (conducted by a CDC-accredited lab) were treated as the gold standard, and pre-vaccination HAI titers from other years were compared against 2015 values for the same individuals. Years with poor correlation (Pearson’s r < 0.5; e.g., 2007, 2008, 2011, 2012) were excluded from downstream analyses. The cohort size is provided in Fig. 1b.

#### Cytokine Measurement

Cytokine levels were measured using Luminex-based multiplex liquid array assays at the Stanford HIMC. All samples were run in duplicate, and control beads (Radix Biosolutions, Georgetwon, TX) were included in every well. Depending on the year and platform version, the following kits were used:

- EMD Millipore Human 80-plex kits (these included 3 panels: Panel 1 was Milliplex HCYTA-60K-PX48; Panel 2 was Milliplex HCP2MAG-62K-PX23; Panel 3 includes the Milliplex HSP1MAG-63K-06 and HADCYMAG-61K-03 (Resistin, Leptin and HGF) to generate a 9 plex): Samples were diluted 3× for Panels 1 & 2 or 10× for Panel 3, with 25 µL of sample used per well. After overnight incubation with antibody-linked magnetic beads at 4°C with shaking, plates were washed twice with wash buffer in a BioTek ELx405 washer (BioTek Instruments, Winooski, VT). Cytokines were detected with biotinylated secondary antibodies and streptavidin-phycoerythrin (PE) (incubations of 1 h and 30 min, respectively, at room temperature with shaking). Plates were washed again as above. The readout was performed on a Luminex FlexMap3D instrument, with a minimum bead count threshold of 50 per cytokine.
- EMD Millipore Magnetic Bead kits (62- or 63-plex): Similar protocol as above, with overnight sample incubation and FlexMap3D readout. Wells yielding bead counts <50 for a given analyte were flagged for quality control.
- Affymetrix/eBioscience Polystyrene Bead kits (37-, 50-, or 51-plex): Samples were incubated on filter-bottom plates (room temperature followed by 4 °C), then washed and detected as per kit instructions. Plates were read on a Luminex 200 instrument, requiring at least 100 beads per cytokine for data inclusion.

#### Antibody Response Quantification

#### HAI titers were measured from serum collected on Day 0 (pre-vaccination) and Day 28 (post-vaccination, as previously described ^1^. Serially diluted sera (25 µL in PBS) were mixed with 25 µL of virus containing 4 HA units in V-bottom 96-well plates. After 15 minutes at room temperature, 50 µL of 0.5% chicken red blood cells (cRBCs) were added, followed by 1-hour incubation. The HAI titer was defined as the reciprocal of the highest serum dilution that inhibited hemagglutination, indicated by a compact cRBC pellet. For each individual, the antibody response was defined as the geometric mean fold-rise in titer across vaccine strains, calculated as the ratio of the Day 28 titer to the Day 0 titer. Fold-rise values were log₂-transformed to normalize distributions. To allow comparison across study years, titers were further adjusted using quantile normalization. Subjects with baseline titers ≥40 were optionally excluded in some analyses to minimize the effects of pre-existing immunity.

#### Multi-cohort Correlation and Meta-analysis

#### Subjects were stratified into subpopulations based on age group, gender, vaccine dose (standard vs. high), baseline HAI titer quantile, study cohort, and influenza season. Each subpopulation (n ≥ 5) was analyzed using linear regression to determine the relationship between the standardized pre-vaccination cytokine level and the log₂ antibody response. The resulting regression slope (with its standard error) was taken as the subgroup's cytokine–response effect size. We then performed a meta-analysis to synthesize these effects across all subgroups, using the rmeta package in R to fit a random-effects model. This approach provided an overall estimate of the correlation between pre-vaccine cytokine levels and antibody response while accounting for between-group heterogeneity.

#### Standardization of Cytokine Concentrations

We standardized cytokine abundances using z-score normalization, allowing us to plot them across different years (Fig. S7). Specifically, we computed a z-score for each cytokine using the mean and standard deviation derived from a reference subset of relatively young subjects (<40 years old) across all years. We selected this younger group, assuming they have more stable immune profiles over time, providing a consistent baseline. This approach is analogous to previously published methods for longitudinal immune monitoring ^2^.

#### Disease Correlation Analysis

#### We investigated associations between baseline cytokine levels and subsequent disease diagnoses in older adults. Seniors (>65 years) from Flu Vaccine Study No. 15 reported new disease diagnoses in annual medical surveys. For each disease of interest, we compared the pre-vaccination IFN-β levels in samples collected within 2 years before diagnosis (cases) with those from disease-free, age-matched subjects (controls). Only cytokine data from subjects >65 years were used in this analysis. Group differences in IFN-β were evaluated to identify any pre-diagnostic cytokine elevations.

#### Spleen Organoid Culture and Vaccination

Spleen tissues were retrieved from the Donor Network West (DNW, an authorized organ processing organization).  The procedure is approved by Stanford IRB (Exemption as donors are deceased).  Spleen and tonsil organoid cultures were established using previously described methods ^3^. Tissue was dissected into roughly 3-5mm and manually disrupted into a single-cell suspension by processing through a 100-μm strainer with a 5mL syringe plunger. Enzymatic dissociation was unnecessary and did not improve the response to LAIV from cryopreserved cells. Tissue debris was reduced by Ficoll density gradient separation, although this step was not required for tonsil organoid development. After washing with complete medium (RPMI with glutamax (ThermoFisher # 61870127), 10% FBS, 1× nonessential amino acids (ThermoFisher #11140050), 1× sodium pyruvate (ThermoFisher # 11360070), 1× penicillin–streptomycin (ThermoFisher # 15240062), 1× Normocin (InvivoGen), and 1× insulin/selenium/transferrin cocktail ( ThermoFisher # 41400045) (Gibco), cells were enumerated and frozen into aliquots in FBS + 10% DMSO. Frozen cells were stored at −140 °C until use.

Aliquots were thawed into complete medium for culture of cryopreserved cells, enumerated, and resuspended to 6 × 107 cells per ml for larger cultures or 2 × 107 cells per ml for smaller cultures. Cells were plated, 100 μl per well, into permeable (0.4-μm pore size) membranes (24-well size PTFE or polycarbonate membranes in standard 12-well plates or 96-well polycarbonate membrane plates with single-well receiver trays; Corning or Millipore), with the lower chamber consisting of complete medium (1 ml for 12-well plates, 200 μl for 96-well plates) supplemented with 1 μg/mL−1 of recombinant human B cell-activating factor (BAFF; BioLegend # 559608). Adding a small amount of BAFF improved total B cell survival (and thus increased overall cell recovery) but was not a requirement for plasmablast differentiation or antibody secretion. Vaccination was performed using Fluzone (IIV, diluted 1:10,000) or FluMist (LAIV, diluted 1:2,000), dose-optimized for organoid stimulation.

#### Cytokine and Antibody Detection in Organoids

#### Cytokine Quantification - Cytokine abundance in organoid culture supernatants was measured using a DNA-barcoded, nanoparticle-based ultrasensitive immunoassay (NULISA, Alamar Biosciences), capable of femtomolar-level detection. Supernatants were collected at Day 3 post-stimulation for cytokine profiling. Each sample (25 µL) was analyzed on the NULISA Inflammation Panel targeting approximately 250 cytokines and chemokines, using the automated ARGO HT platform at the Stanford Human Immune Monitoring Center (HIMC). For stimulation studies comparing IIV, LAIV, IFN-β, and combinations thereof, culture conditions were standardized, and cytokine abundance was reported relative to unstimulated controls matched by donor.

#### Influenza-A-specific IgG Detection - To evaluate antigen-specific antibody responses, influenza hemagglutinin (HA)-specific IgG was quantified in organoid supernatants at Day 7 post-vaccination using a Luminex-based multiplex immunoassay. For the cytokine adjuvant screen, 19 recombinant human cytokines were co-administered with inactivated influenza vaccine (IIV) at three concentrations (1, 10, or 100 ng/mL, with adjusted ranges for IL-1β and IL-18), and influenza-specific IgG was measured relative to IIV-only controls. Organoids were generated from dissociated human spleen and tonsil tissue and maintained in a 96-well format for high-throughput screening. All measurements were performed in biological replicates (n = 5 donors), and fluorescence intensities were normalized and log-transformed to assess fold changes over background stimulation levels.

#### Low-Cell-Input Organoid Culture

Organoids were established using low-input cell seeding to scale up culture throughput. Cells were seeded in ultra-low attachment 96-well plates at 1.6 × 10⁵ cells/well in 200 µL of culture media. Media was partially exchanged (30%) every two days. On Day 7, culture supernatants were harvested for antibody quantification as described above.

#### NULISAseq Inflammation Panel

The NULISAseq Inflammation Panel (Alamar Biosciences, Fremont, California), performed at the Stanford University HIMC, is a multiplexed proximity ligation assay targeting 250 inflammation-associated proteins. The assay was processed automatically in the ARGO HT system (Alamar BioSciences). 25 μl of each sample was loaded on the sample plate, along with three sample controls (SC), four negative controls (NC), and three inter-plate controls (IPC). After completion of the automated run, next-generation sequencing (Illumina, Foster City, California) was performed on the pooled library. Data were generated using ACC (Alamar Command Center) and NAS (NULISA Analysis Software) via normalization to Internal Controls (IC) and Inter-Plate Controls (IPC). Raw counts were normalized to internal and inter-plate controls and then log₂-transformed to yield normalized protein quantity (NPQ) values.

#### Viral Antibody Assay

As previously described, a custom Luminex assay was created to detect antibody responses to SARS-CoV-2 and other viral antigens ^4^. Antigens of interest were coupled to barcoded beads (Luminex Corporation, Austin, Texas) according to the manufacturer’s instructions. Supernatant samples were run undiluted, with 25 μl of sample added to the assay plate containing the antigen-coupled bead mixture and incubated for 2 hours at room temperature or overnight at 4°C, shaking on an orbital shaker. Samples were then washed, and 25 μl of secondary Goat-anti Human IgG (Fc fragment) coupled to PE (Phycoerythrin) (Anti IgG-Cat# NC9822979, Jackson ImmunoResearch, West Grove, PA) was added. After incubation with shaking for 30 minutes at room temperature, a second wash was performed before adding 130 μl Reading Buffer (Luminex). Samples were read on a Luminex Flex 3D instrument with a lower bound of 50 beads per target antigen. This assay was performed by the Stanford University HIMC and is presented as MFI (Median Fluorescence Intensity).

**RNA Vaccine Synthesis Protocol**

DNA sequences for vaccines were codon optimized using an in-house algorithm and were synthesized by Synbio Technologies (Monmouth Junction, NJ). 3’ and 5’ UTRs are taken from the commercial Pfizer vaccine ^5^. Gene sequences were PCR amplified and cloned into a backbone containing 3’ and 5’ UTRs (NEBuilder, NEB). Then, constructs were amplified, linearized with a T7 promoter 5’ overhang, and polyadenylated via a 2-step PCR. PCR Steps were performed using 0.5 µm each primer, 1-2 ng template per 50 µL reaction, and Platinum SuperFi II polymerase (Thermo Fisher). PCRs were cleaned up via standard protocol (Qiagen). In vitro transcription of purified PCR products was performed with the T7 mScript Standard mRNA production kit (Cellscript) while using N1-methylpseudouridine (Trilink Biotech) instead of uracil. Then, RNA was capped using the ScriptCap Cap 1 Capping System (Cellscript). RNA was purified via the Monarch RNA Cleanup kit. IVT RNA was stored at -80 °C. The following lipid solution is prepared to make the LNPs. The lipids are the same as those used in the commercial Pfizer vaccine ^5^.

| Reagent | Working Conc. (mg/mL) | Working Conc. (mM) | Stock Conc. | Volume per 1mL Stock (µL) |
| --- | --- | --- | --- | --- |
| ALC-0315 | 5.41 | 7.06 | 50mg/mL | 108.2 |
| 100% Ethanol | N/A | N/A | N/A | 150.9 |
| ALC-0159 | 0.629 | 0.26 | 10mg/mL | 62.9 |
| Cholesterol | 2.52 | 6.52 | 10mg/mL | 252 |
| DSPC | 1.13 | 1.43 | 5mg/mL | 226 |
| 50% Ethanol | N/A | N/A | N/A | 200 |

RNA is thawed and diluted to final concentrations of 141ng/uL RNA, 25mM Acetate, 200mM NaCl, pH 4. Then, RNA and lipid solutions undergo microfluidic mixing at a 3 volume ratio of RNA: 1 volume lipid. Solutions are loaded into 1 mL syringes, as larger plastic syringes flex under pressure and are unsuitable for mixing. It is essential that the syringes contain nearly no air pockets, or mixing will not occur at a reliable rate. For small batches, syringes can be pre-filled with buffer, and a small air pocket can be used to separate RNA from RNA buffer along the length of the tubing. For large batches, the 1 mL syringes are filled. Solutions flow from the syringe to the microfluidic mixer via Cole Parmer Masterflex Microbore Transfer Tubing (Tygon® ND-100-80, 0.020" ID x 0.060" OD), and into the device via custom tubing adapters (0.025” OD, 0.013” ID, 0.5” length type 304 stainless steel, New England Small Tube). The overall flow rate is 500 μL per minute through the device, controlled via syringe pump (New Era Pump Systems Inc. NE-4000).

The output solution is dialyzed against PBS to remove the ethanol (Pierce microdialysis devices, 0.3mL capacity, 3.5 kDa MWCO), typically diluting the final vaccine to 30ug RNA / 400uL. If the LNPs are to be frozen, the vaccine is further diluted by adding 1/2 volume sterile-filtered 60% sucrose in PBS (final sucrose is 20% w/v). LNPs are then flash frozen in liquid nitrogen and placed in LN2 storage. The microfluidic mixer used for LNP synthesis was fabricated using standard photolithography; the corresponding photomask design is available upon request. Photomasks were ordered from Artnet Pro Inc. (San Jose, CA). The flow channel layer was fabricated to a height of roughly 63 microns, and the herringbone layer was manufactured to a height of approximately 21 microns. The photomask was drafted in-house, but the basis for this mixer was previously published ^6^.

#### Mice and Immunization

Male C57BL/6J mice (6-10 weeks old) were obtained from the Jackson Laboratory and used for all experiments described in this study. Mouse were housed in the Stanford Animal Facility under specific pathogen-free (SPF) conditions, maintained on a 12-hour light/12-hour dark cycle at a temperature of ~18-23 °C and 40–60% humidity. All animal procedures were reviewed and approved by the University Administrative Panel on Laboratory Animal Care (APLAC; protocol No. 34513).

For immunization, each mouse received a total of 120μl of intramuscular injections, administered as 60μl into each hind leg (left and right caudal thigh muscles). The injection mixture contained 2023/24 inactivated influenza vaccine (IIV; Fluzone® Quadrivalent) at a dose of 1 or 10μg per mouse, combined with lipid nanoparticle (LNP)-encapsulated mRNA encoding mouse cytokine adjuvants at 1 or 3μg per mouse.

#### Fluorescence imaging

To assess the distribution of LNP-encapsulated mRNA transcripts, mice were intramuscularly immunized with PBS or LNP-encapsulated mRNA encoding GFP (3μg per mouse) as described. Draining lymph nodes and hind limb muscles were harvested at 24 hours post-immunization. Whole-tissue fluorescence was then measured using a Largo spectral imaging system with an excitation wavelength of 465 nm and an emission wavelength of 510 nm. The data were analyzed using Aura imaging software (Spectral Instruments Imaging) and values represent the integrated fluorescence intensity.

| **Table S1: Literature Summary of Cytokine Adjuvant Effect (IgG production) in Mouse Studies** | | | | | | |
| --- | --- | --- | --- | --- | --- | --- |
| **Cytokine Tested** | **CytokineFormat** | **Antigen Species** | **Antigen** | **Delivery Route** | **Effect on IgG** | **Ref** |
| IFNα/β | Protein | None - Model Antigen | Chicken Gamma Globulin | SC | Pos | ^7^ |
| IFNα/β | Protein | Influenza | Influenza Vaccine | IM | Pos | ^8^ |
| IFNγ | Protein | HIV | gp-120 | IP | Pos | ^9^ |
| IFNγ | Protein | Influenza | Influenza Vaccine | IP | Pos | ^10^ |
| IL-10 | DNA | HIV | Gag/pol; ENV | IM | Pos | ^11^ |
| IL-12 | Protein | Influenza | H1, N1 | IN | Pos | ^12^ |
| IL-15 | Protein | *Staphylococcus aureus* | STEBVax toxoid | IM | Pos | ^13^ |
| IL-17 | Protein | BoHV-5 | BoHV-5 gD | IM | Pos | ^14^ |
| IL-18 | DNA | HIV | Gag/pol; ENV | IM | Pos | ^11^ |
| IL-18 | DNA | *Schistosoma japonicum* | GST | IM | None | ^15^ |
| IL-18 | Protein | Influenza | HA | IN | Pos | ^16^ |
| IL-18 | RNA | Influenza | Split Influenza Vaccine | IM | Pos | ^17^ |
| IL-1α/β | Protein | None - Model Protein | BSA | IP | Pos | ^18^ |
| IL-1α/β | Protein | None - Model Antigen | Ovalbumin | MOP | Pos | ^19^ |
| IL-1α/β | Protein | Influenza | HA | IN | Pos | ^16^ |
| IL-1α/β | Protein | None - Model Antigen | Ovalbumin or TT | IN | Pos | ^20^ |
| IL-1β | Protein | *Clostridium difficile* | Flagellar Cap FliD Protein | OG | None | ^21^ |
| IL-2 | DNA | HIV | Gag/pol; ENV | IM | Pos | ^11^ |
| IL-2 | DNA | Influenza | Influenza Vaccine | SC | Pos | ^22^ |
| IL-21 | DNA | HIV | ENV Peptide | IM | Neg | ^23^ |
| IL-21 | DNA | HBV | HBV Envelope | IM | None | ^24^ |
| IL-21 | DNA | None - Tumor Model | SCCVII | IV | Pos | ^25^ |
| IL-21 | DNA | None- Tumor Model | GD2 | IM | Pos | ^26^ |
| IL-21 | DNA | HIV | Gag + MVA | IM | None | ^27^ |
| IL-21 | Cell | TB | ESAT-6 | SC | None | ^28^ |
| IL-4 | DNA | HIV | Gag/pol; ENV | IM | Pos | ^11^ |
| IL-4 | DNA | Influenza | Influenza Vaccine | SC | Pos | ^22^ |
| IL-5 | DNA | HIV | Gag/pol; ENV | IM | Pos | ^11^ |
| IL-9 | DNA | FMDV | VP1 | IM | Pos | ^29^ |
| LTA | DNA | HIV | Gag/pol; ENV | IM | Pos | ^11^ |
| TNF | DNA | HIV | Gag/pol; ENV | IM | Pos | ^11^ |

Abbreviations: Subcutaneous (SC), Intramuscular (IM), Intraperitoneal (IP), Intranasal (IN) Mini Osomotic Pump (MOP), Intravenous (IV). Effect on IgG determined by author description or based on average IgG fold change or IgG O.D. change values when provided Pos (>1X with strong trend or significance), None (1X, inconsistent, or not significant), Neg (<1X).
